# Metabolite Quantitative Trait Loci for flavonoids provide new insights into the genetic architecture of strawberry (*Fragaria* x *ananassa*) fruit quality

**DOI:** 10.1101/2020.03.20.000455

**Authors:** Marc Labadie, Guillaume Vallin, Aurélie Petit, Ludwig Ring, Thomas Hoffmann, Amèlia Gaston, Aline Potier, Wilfried Schwab, Christophe Rothan, Béatrice Denoyes

## Abstract

Flavonoids are products from specialized metabolism that contribute to fruit sensorial (colour) and nutritional (antioxidant properties) quality. Here, using a pseudo full-sibling F_1_ progeny previously studied for fruit sensorial quality of cultivated strawberry (*Fragaria* x *ananassa*), we explored over two successive years the genetic architecture of flavonoid-related traits using LC-ESI-MS (13 compounds including anthocyanins, flavonols and flavan-3-ols) and colorimetric assays (anthocyanins, flavonoids, phenolics, FRAP and TEAC antioxidant capacity). Network correlation analysis highlighted the high connectivity of flavonoid compounds within each chemical class and low correlation with colorimetric traits except anthocyanins. Mapping onto the female and male linkage maps of 152 flavonoid metabolic QTLs (mQTLs) and of 26 colorimetric QTLs indicated co-localization on few linkage groups of major flavonoid- and taste-related QTLs previously uncovered. These results pave the way for the discovery of genetic variations underlying flavonoid mQTLs and for marker-assisted selection of strawberry varieties with improved sensorial and nutritional quality.

## Introduction

Consumers are more and more aware that consumption of fruits and vegetables has long-term effects on human health.(1,2,3). In this respect, small fruits including strawberry are an important source of phytochemicals with proven health-benefits.(4–7). The most consumed small fruit worldwide is cultivated strawberry (*Fragaria* x *ananassa*), which is issued from the hybridization few centuries ago between *F. chiloensis* and *F. virginiana*.(8). Since then, cultivated strawberry has undergone intense breeding activity for traits related to yield and cultural practices (9) but also for sensorial fruit quality traits such as fruit weight and shape, shininess and colour and, more recently, flavor.(10). In the last decades, with the increased interest in nutritionally active phytochemicals from fruits, major strawberry fruit antioxidants such as ascorbic acid (vitamin C), ellagic acid and other polyphenolic compounds have been studied more deeply.(3,5,6,11).

In contrast to ascorbate, for which QTLs have recently been mapped and candidate genes have been identified (11), the genetic architecture of polyphenol-dependent nutritional traits remains poorly known in cultivated strawberry. To date, detailed analyses of flavonoid composition have been performed on both cultivated and wild diploid *Fragaria vesca* strawberry species.(12–14). In addition, major strawberry flavonoid genes have been isolated and the activity of a few corresponding enzymes have been characterized.(15–17). In the wild diploid strawberry species *Fragaria nilgerrensis*, a natural mutation in the MYB10 transcription factor regulating anthocyanin biosynthesis is responsible for the white fruit phenotype.(18). However, with a few exceptions such as the mapping of a peroxidase that controls the trade-off between lignin and anthocyanins biosynthesis to a locus responsible for fruit color variations (19), the extent to which natural genetic variations may control flavonoid content in cultivated strawberry remains poorly known. Such studies are further complicated by the octoploid status of cultivated strawberry (2n = 8x = 56) in which variations in a given trait at a single locus can be controlled by up to eight homoeoalleles located on four linkage groups (20) corresponding to four different subgenomes (*F. iinumae*, *F. nipponica*, *F. viridis* and *F. vesca*).(21).

To get more insights into the genetic control of flavonoid content, the first step is to isolate and identify the most abundant flavonoids present in strawberry fruit and then to map the corresponding metabolic Quantitative Trait Loci (mQTL) onto cultivated strawberry genetic map. State-of-the art methods now allow the exhaustive analysis of flavonoids, which are derived from the phenylpropanoid pathway.(22). Flavonoids (anthocyanins, flavonols, flavan-3-ols) found in strawberry fruit are antioxidant molecules that have proven dietary health-benefits.(2,23,24). Flavonols in cultivated strawberry are mainly glycosides of quercetin and kaempferol. Most common flavan-3-ols are catechin and epicatechin as well as their monohydroxylated equivalents afzlechin and epiafzlechin and glycosylated derivatives. Anthocyanins are mainly glycosides of pelargonidin and cyanidin.(25). In addition to their antioxidant properties, anthocyanins are water-soluble pigments that give to the strawberry fruit its attractive bright red (pelargonidin derivatives) to dark red (cyanidin derivatives) colour. Pelargonidin derivatives are the most abundant anthocyanins in cultivated strawberry (19, 23) while pelargonidin and cyanidin derivatives are equally found in *F. vesca*.(14). Although several colour-related QTLs have been detected (20), the genomic regions responsible for quantitative variations of the various flavonoid compounds found in cultivated strawberry fruit have not been uncovered to date.

In this study, to further explore the genetic architecture of polyphenolic- and flavonoid-dependent traits in cultivated strawberry, we analyzed over two successive years (with three repeats each year) a pseudo full-sibling F_1_ progeny previously studied for the genetic control of fruit sensorial quality.(20). The population also segregates for flavonoids (13 compounds from three chemical classes) and for colour- and antioxidant-related traits (5 traits assayed by colorimetry). In total, we detected 152 mQTLs for the flavonoid compounds, all of which were identified for the first time from cultivated strawberry, and 26 QTLs for the colour- and antioxidant-related traits. The largest number of flavonoid mQTLs and of colour- and antioxidant-related QTLs was detected on the LG VIa linkage group that also harbors major fruit sweetness and acidity-related traits (20) and is therefore a likely target for the improvement by Marker-Assisted Selection (MAS) of the sensorial and nutritional quality of strawberry fruit.

## Material and methods

### Plant materials and preparation

A pseudo full-sibling F_1_ population of 165 individuals obtained from a cross between the variety ‘Capitola’ (CA75.121-101 x Parker, University of California, Davis, USA) and the advanced line ‘CF1116’ ([‘Pajaro’ x (‘Earlyglow’ x ‘Chandler’)], reference from Ciref, France) was developed. The two parents, ‘Capitola’ and ‘CF1116’, display many contrasting fruit quality traits.(20, 26). For each of the two consecutive study years (2010 and 2011), six cold-stored strawberry plants per genotype, which were planted in 2009 and 2010, were grown in soil-free pine bark substrate under plastic tunnel with daily ferti-irrigation and control of biotic stresses. Mapping population included a total of 165 individuals over the two study years.

Within this progeny, 72 and 131 individuals including parents were respectively phenotyped in 2010 and in 2011. Fruits were harvested at red ripe stage, when red coloration of the fruit is homogeneous. To ensure conformity of the fruit samples and to avoid undesirable effects on antioxidant and fruit polyphenolic contents, several precautions were taken. Fruits were collected in the morning and those showing abnormal shape and size, or injuries were not harvested. For each genotype, two harvests of 4-8 fruits each were performed at the peak of fruit production (8-16 fruits in total). Fruits were immediately frozen in liquid nitrogen and stored at −80°C, in order to avoid degradation of antioxidants and polyphenolics. They were then ground into a fine powder in liquid nitrogen and frozen powders from two successive harvests were pooled (volume/volume) and blended. Three samples of pooled frozen powder were used for the various extractions.

### Antioxidant measurements

Extraction of hydrophilic antioxidants was adapted from Capocasa et al. (27). For each sample, 0.7 +/− 0.02 g of fruit powder was dissolved in the dark in 7 ml of methanol/water (80/20 v/v) extraction solution. The mixture was then vortexed for 30 s, agitated at 160 rpm for 30 min and centrifuged for 10 min at 4,500 g. All steps were performed at ∼5°C in the dark. Aliquotes of 250 µl of supernatant (fruit extract) were then stored at −80°C in microtubes until colorimetric analyses.

Five antioxidant-related traits were measured on fruit extracts by colorimetric assays including anthocyanin content (ANTHc), flavonoid content (FLAVc), total phenolic content (PHENc) and the two antioxidant-related traits FRAP (Ferric Reducing Antioxidant Power) and TEAC (Trolox Equivalent Antioxidant Capacity). ANTHc was estimated according to the protocol of Villarreal et al. (28). Results are expressed as mg pelargonidin-3-glucoside equivalents/100 g fresh weight. FLAVc was estimated by the method of Dewanto et al. (29). Results are expressed as µg catechin equivalents/g fresh weight. PHENc was estimated by the method of Sinklard and Singleton.(30). Results are expressed as mg gallic acid equivalents/g fresh weight. FRAP and TEAC were determined according to Benzie and Strain (31) and Re et al. (32), respectively. Results are expressed as µm Trolox equivalents/g fresh weight for both FRAP and TEAC. For each genotype and colorimetric assay, 4 technical repeats from the pooled two-harvest-fruit-powder were performed except for FRAP in 2010 (2 technical repeats).

### LC–ESI–MS^n^ analysis of polyphenolic metabolites

A total of 13 individual phenolic metabolites [anthocyanins (pelargonidin-3-glucoside; pelargonidin-3-glucoside-malonate; pelargonidin-3-rutinoside; cyanidin-3-glucoside (kuromanin); (epi)afzelechin-pelargonidin-glucoside), flavonols (kaempferol-glucoside; kaempferol-glucuronide; kaempferol-cumaroyl-glucoside; quercetin-glucuronide), flavan-3-ols (catechin; (Epi)catechin dimers; (epi)afzelechin-(epi)catechin dimers; (epi)afzelechin-glucoside)] were measured by LC–ESI–MS for the two years.

Extraction and LC–ESI–MS*^n^* analysis of polyphenolic metabolites were done as described in Ring et al. (19). Briefly, 500 mg of frozen powder was extracted twice with methanol from each biological replicate, adding biochanin A as internal standard. After centrifugation, the supernatants were combined, dried under vacuum, dissolved in water and the samples were injected twice as a technical replicate. Samples were analysed on an Agilent 1100 HPLC/UV system (Agilent Technologies, Waldbronn, Germany) equipped with a reversed phase column (Luna® 3 μm C18(2) 100Å 150 × 2 mm, Phenomenex, Aschaffenburg, Germany) and connected to a Bruker Esquire 3000 plus ion trap mass spectrometer (Bruker Daltonics, Bremen, Germany). Data were analysed with Data analysis 5.1 software (Bruker Daltonics, Bremen, Germany). Metabolites were identified by comparing their retention times and mass spectra (MS and MS2) with those of measured authentic reference compounds. The major known phenolic metabolites were quantified in the positive and negative MS mode by the internal standard method, using QuantAnalysis 2.0 (Bruker Daltonics, Bremen, Germany). Results are expressed as mg equivalents /100 g fresh weight assuming a response factor of 1. Analyses of pooled frozen powder samples were carried out in 2010 and 2011 on 6 replicates for parents and on 3 replicates for individuals from the progeny.

### Data analysis

Exploratory analyses were performed on all individuals (72 in 2010 and 131 in 2011) and parents (‘Capitola’ and ‘CF1116’) using R software (R 3.5.0) in the interface RStudio (RStudio 1.2.1572). A Kruskal-Wallis test (ANOVA on the rank and appropriate post hoc test) was used to compare the mean values between the parents (agricolae 1.3.1 and PMCMR 4.3 R packages). Trait segregation was declared transgressive when at least one progeny had a value that was higher or lower than that of the highest or lowest parent, by at least twice the standard deviation of the parents.(20). Phenotypic correlations were estimated as Pearson correlations between each trait and represented by heatmap using stats 3.5.0 and corrplot 0.84 R packages and by network correlation using ggnet 0.1.0 and network 1.15 R packages (correlation values r > 0.3 for highlighting the strong correlations). Where a genotypic effect within progeny was found, broad sense heritability was evaluated from variance analysis as follows:

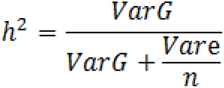

With VarG: Genetic variance, Var*e*: residuals’ variance, n= number of repetitions (in our case n=3). VarG and Var*e* were determined using ANOVA on linear regression of each trait.

### QTL detection

To allow comparison with previously published results, QTL detection and analysis were performed as described in Lerceteau-Köhler et al. (20). Briefly, QTL detection was performed by composite interval mapping (CIM) (33, 34) using model 6 of QTL Cartographer software, in which five co-factors were included. The statistical significance thresholds (LOD value) for both female and male linkage maps, and for declaring a putative QTL were 2.8 or 3.1 after 1.000 permutation times and significance level of α = 0.10 or 0.05 respectively. The principal characteristics of mQTLs and colour- and antioxidant-related traits’ QTL included the chromosome, marker, position, LOD value, confidence interval (LODmax ± 1 LOD), and the proportion in % of phenotypic variance (R^2^) explained by a single QTL. When QTLs from two or three replicates of a same year overlapped, they were summarized in a single QTL.

## Results

### Flavonoid and antioxidant- and colour-related traits in the ‘Capitola’ and ‘CF1116’ parents

Flavonoid compounds were identified by LC–ESI-MS in the cultivated variety ‘Capitola’ (female), the breeding genotype ‘CF1116’ (male) and in the pseudo full-sibling F_1_ progeny issued from a cross between them. The two parents are contrasted for several fruit quality traits including fruit shape and weight, firmness and fruit sweetness, acidity and colour-related traits. (20). Among the putative metabolites observed, 13 flavonoid compounds were unambiguously identified in 2010 and 2011 by comparison with commercial standard run under the same conditions. The identified compounds included five anthocyanins (pelargonidin-3-glucoside, pelargonidin-3-glucoside-malonate, pelargonidin-3-rutinoside, cyanidin-3-glucoside and (epi)afzelechin-pelargonidin-glucoside), four flavonols (kaempferol-glucoside, kaempferol-glucuronide, kaempferol-coumaryl-glucoside and quercetin-glucuronide) and four flavan-3-ols (catechin, (epi)catechin dimer, (epi)afzelechin-(epi)catechin dimer and (epi)afzelechin-glucoside). Corresponding abbreviations are indicated in Table 1. The metabolic compounds identified were further summed up by chemical family to give total anthocyanins (Ant), total flavonols (Fvo) and total flavan-3-ols (F3ol). We also analyzed five traits by colorimetric assays: anthocyanin content (ANTHc), flavonoid content (FLAVc), phenolic content (PHENc) and antioxidants (FRAP and TEAC). In 2011, the FRAP trait was not measured. The general distribution parameters of the metabolites and of the additional antioxidant- and colour-related traits were evaluated for the two parents (‘Capitola’ and ‘CF1116’) and for the progeny in 2010, and for only ‘Capitola’ and for the progeny in 2011. In order to facilitate comparison of the progeny with the two parents, the results of metabolite profiling and colorimetric assays for 2010 are shown in Table 1. Results for 2011 are provided as Supplemental Table S1.

**Table 1.** Trait values for ‘Capitola’ and ‘CF1116’ parents and related progeny in 2010.

| Traits | Abbr. | ‘Capitola’ |  |  | ‘CF1116’ |  |  |  | Progeny |  |  | *h*<sup>2</sup> | Tr |
| --- | --- | --- | --- | --- | --- | --- | --- | --- | --- | --- | --- | --- | --- |
|  |  | mean | s.d. | range | mean | s.d. | Range | Cap vs CF | mean | s.d. | range |  |  |
| Anthocyanins |  |  |  |  |  |  |  |  |  |  |  |  |  |
| Total anthocyanins | Ant | 26.88 | 6.81 | 19.37-32.63 | 26.62 | 4.29 | 21.76-29.84 | n.s. | 24.18 | 6.94 | 10.55-57.64 | 0.77 | + & - |
| Pelargonidin-3-glucoside | PgGs | 24.15 | 5.40 | 18.00-28.15 | 25.17 | 3.92 | 20.71-28.04 | n.s. | 22.37 | 6.32 | 10.19-53.47 | 0.81 | + & - |
| Pelargonidin-3-glucoside-malonate | PgGsM | 0.19 | 0.09 | 0.10-0.26 | 0.31 | 0.14 | 0.21-0.47 | n.s. | 0.17 | 0.30 | 0.00-2.12 | 0.93 | + & - |
| Pelargonidin-3-rutinoside | PgRs | 2.35 | 1.47 | 1.13-3.99 | 0.64 | 0.15 | 0.47-0.74 | * | 1.25 | 0.89 | 0.04-4.56 | - | - |
| Cyanidin-3-glucoside | CyGs | 0.18 | 0.04 | 0.14-0.23 | 0.48 | 0.12 | 0.35-0.57 | * | 0.36 | 0.24 | 0.07-1.63 | 0.92 | + & - |
| (epi)Afzelechin-pelargonidin-glucoside | AfPgGs | 0.001 | 0.002 | 0.000-0.003 | 0.017 | 0.002 | 0.015-0.019 | * | 0.01 | 0.01 | 0.00-0.03 | 0.57 | + |
| Flavonols |  |  |  |  |  |  |  |  |  |  |  |  |  |
| Total Flavonols | Fvo | 0.24 | 0.05 | 0.18-0.28 | 0.32 | 0.04 | 0.28-0.34 | n.s. | 0.30 | 0.09 | 0.13-0.73 | 0.94 | + & - |
| Kaempferol-glucoside | KGs | 0.09 | 0.03 | 0.06-0.12 | 0.14 | 0.01 | 0.12-0.15 | * | 0.10 | 0.04 | 0.03-0.22 | 0.96 | + & - |
| Kaempferol-glucuronide | KGn | 0.05 | 0.01 | 0.04-0.06 | 0.045 | 0.003 | 0.04-0.05 | n.s. | 0.06 | 0.02 | 0.02-0.15 | 0.88 | + & - |
| Kaempferol-coumaryl-glucoside | KCoGs | 0.07 | 0.01 | 0.05-0.08 | 0.09 | 0.01 | 0.08-0.10 | n.s. | 0.09 | 0.03 | 0.03-0.19 | 0.92 | + & - |
| Quercetin-glucuronide | QGn | 0.03 | 0.01 | 0.02-0.04 | 0.05 | 0.01 | 0.04-0.07 | * | 0.05 | 0.04 | 0.02-0.27 | 0.98 | + & - |
| Flavan-3-ols |  |  |  |  |  |  |  |  |  |  |  |  |  |
| Total Flavan-3-ols | F3ol | 1.07 | 0.23 | 0.81-1.22 | 0.82 | 0.07 | 0.75-0.90 | n.s. | 1.21 | 0.36 | 0.58-2.70 | 0.94 | + & - |
| Catechin | Cat | 0.19 | 0.04 | 0.15-0.23 | 0.20 | 0.02 | 0.19-0.23 | n.s. | 0.22 | 0.05 | 0.10-0.42 | 0.78 | + & - |
| (epi)Catechin dimers | CatCat | 0.48 | 0.13 | 0.34-0.56 | 0.47 | 0.06 | 0.40-0.52 | n.s. | 0.47 | 0.11 | 0.21-0.97 | 0.85 | + & - |
| (epi)Afzelechin-(epi)catechin dimers | AfCat | 0.04 | 0.01 | 0.03-0.04 | 0.05 | 0.01 | 0.04-0.06 | * | 0.05 | 0.02 | 0.01-0.13 | 0.81 | + & - |
| (epi)Afzelechin-glucoside | AfGs | 0.36 | 0.09 | 0.29-0.46 | 0.11 | 0.03 | 0.08-0.14 | * | 0.47 | 0.31 | 0.11-1.95 | 0.98 | + |
| Colorimetry |  |  |  |  |  |  |  |  |  |  |  |  |  |
| Anthocyanins (colorimetry) | ANTHc | 832.82 | 7.63 | 824.5-839.5 | 461.05 | 5.39 | 454.9-464.7 | * | 652.96 | 191.92 | 249.8-1131.4 | 1 | + & - |
| Flavonoids (colorimetry) | FLAVc | 625.62 | 13.39 | 611.2-637.7 | 392.81 | 36.68 | 366.9-418.8 | n.s. | 465.77 | 164.15 | 0.0-788.2 | 0.66 | + & - |
| Phenolics (colorimetry) | PHENc | 1.71 | 0.55 | 1.28-2.33 | 1.12 | 0.56 | 0.66-1.75 | n.s. | 1.37 | 0.48 | 0.54-3.24 | - | + |
| Antioxidant FRAP | FRAP | 7.77 | 1.39 | 6.79-8.75 | 6.76 | 1.48 | 5.71-7.81 | n.s. | 7.59 | 1.44 | 4.60-12.06 | 0.49 | + |
| Antioxidant TEAC | TEAC | 14.37 | 1.21 | 12.98-15.19 | 11.79 | 0.49 | 11.29-12.28 | * | 13.62 | 2.15 | 7.98-19.55 | 0.8 | + & - |

For each trait, the mean value, standard deviation (s.d.), range and heritability (h<sup>2</sup>) are described.
The significance levels of the differences of the means between ‘Capitola’ and ‘CF1116’ are indicated in the Cap vs CF column and were analyzed by a test of
ANOVA-on the rank (Kruskal-Wallis test and post-hoc test). n.s., \*, \*\* and \*\*\* indicate values that are not significant (n.s) or are significant at 0.01 <P< 0.05,
0.001 <P< 0.01, and P <0.001, respectively. Trait segregation was declared transgressive (Tr) when at least one progeny had a value higher (+), lower (-) or both (+ & -) than the highest or lowest parental value by at least twice the s.d. of the parents. Flavonoid metabolites values are expressed as mg-equ/100 g FW assuming a response factor of 1. Colour- and antioxidant-related traits: ANTHc results are expressed as mg pelargonidin-3-glucoside equivalents/100 g FW. FLAVc results are expressed as µg catechin equivalents/g FW. PHENc results are expressed as mg gallic acid equivalents/g FW. FRAP and TEAC results are expressed as µm Trolox equivalents/g FW. Values are the means of n = 3 replicates per genotype.

Mean values were significantly different between the two parents for less than half of the traits analyzed (Table 1). Total flavonoid content, obtained by summation of the anthocyanins, flavonols and flavan-3-ols, was remarkably similar in ‘Capitola’ and ‘CF1116’ (28.2 *vs.* 27.8 mg equ/100 g fresh weight in 2010). Anthocyanins were by far the major contributors (∼93 % of the flavonoids) followed by flavan-3-ols (∼2.9 to 3.7 %). Close examination of the results showed small to large variations for individual compounds within a given chemical family. The content in pelargonidin-3-glucoside, which contributes to the total anthocyanin content for as much as ∼90 %, was similar in both parents. In contrast, pelargonidin-3-rutinoside content (∼2.4 to 8.7 % of the total anthocyanins) was more than 3-fold higher in ‘Capitola’ at the opposite of cyanidin-3-glucoside (∼0.7 to 1.8 % of the total anthocyanins) that was almost 3-fold lower in ‘Capitola’. The main flavonol kaempferol-glucoside (∼37.5 to 43.8 % of the total flavonols) displayed similar values in both parents. Contents in the main flavan-3-ol (epi)catechin dimers (∼44.9 to 57.3 % of the total flavan-3-ols) and in catechin (∼17.8 to 24.4 %) were not much different between the two parents while (epi)afzelechin-glucoside (∼13.4 to 33.6 % of the total flavan-3-ols) was 2.5-fold higher in ‘Capitola’. Surprisingly, given the results from LC-MS analyses that showed similar anthocyanins values for both parents, the colorimetric estimation of anthocyanin content produced a mean ANTHc value 1.8-fold higher in ‘Capitola’ than in ‘CF1116’, in agreement with previously published results.(20). FLAVc, PHENc, FRAP and TEAC mean values were all higher in ‘Capitola’, with FLAVc and PHENc being respectively 1.6-fold and 1.5-fold higher than in ‘CF1116’.

### Distribution of metabolites and antioxidant- and colour-related traits in the ‘Capitola’ x ‘CF1116’ population

Metabolite profiling and measurements of antioxidant- and colour-related traits were next performed on the progeny in order to delineate genomic regions responsible for trait variations. Trait analysis was carried out on 72 individuals in 2010 and 131 genotypes in 2011. Distribution parameters (mean, standard deviation, range and heritability) are shown in Table 1 for 2010. In both years, considerable variations were observed for most of the traits, which segregated in the progeny. Compared to the parents, the range was considerably extended for metabolites belonging to the various chemical classes and for colorimetric traits, thus indicating a genotype-specific control. Most of the variations among the most extreme genotypes were in the 4- to 10-fold range, an example of which is the 5-fold variation for the most abundant compound pelargonidin-3-glucoside. However, the anthocyanins cyanidin-3-glucoside and pelargonidin-3-rutinoside displayed 23-fold and 114-fold variations in the progeny, respectively, while the flavan-3-ol (epi)afzelechin-glucoside exhibited a 17-fold variation. Similar results were reported in 2011 (Supplemental Table S1).

Calculated broad sense heritability displayed high values (h^2^ > 0.5) for all the 21 traits analyzed in 2010 except FRAP (h^2^=0.49). Very high heritability values (from h^2^ > 0.8) were even observed for 14 traits associated with anthocyanins (3), flavonols (5) and flavan-3-ols contents (4) and with ANTH and TEAC (Table 1). Very similar results were obtained in 2011 (Supplemental Table S1). Transgressions were detected in 2010 for all traits. They were both positive and negative for 16 of the 21 traits, but only negative for PgRs and positive for AfPgGs, AfGs and PHENc (Table 1).

### Correlation of metabolites and antioxidant- and colour-related traits in the ‘Capitola’ x ‘CF1116’ population

Pearson phenotypic correlations between the various traits are shown in Figure 1A for 2010 and in Supplemental Figure S1A for 2011. In 2010, Pearson correlation analysis highlighted the strong correlation (r=0.99) between total anthocyanins and PgGs, which is not surprising given that PgGs is the preponderant anthocyanin in cultivated strawberry. It also highlighted significant correlations with minor anthocyanin compounds including PgGsM (r=0.36), PgRs (r=0.48), CyGs (r=0.57) and AfPgGs (r=0.52). In addition, anthocyanin content was positively correlated with that of flavonoids from various chemical classes including flavonols (r=0.30 for QGn to r=0.54 for KGs) and flavan-3-ols (r=0.40 for CatCAt and r=0.38 for AfCat). As could be expected, correlations were also significant between flavonoid compounds belonging to the same chemical classes e.g. between the flavonols KGn and QGn (r=0.53) and the flavan-3-ols CatCat and AfCat (r=0.75). No negative correlation was found except for the weak correlation (r=-0.18) between the anthocyanin PgRs and the flavan-3-ol Cat.

**Figure 1A.**
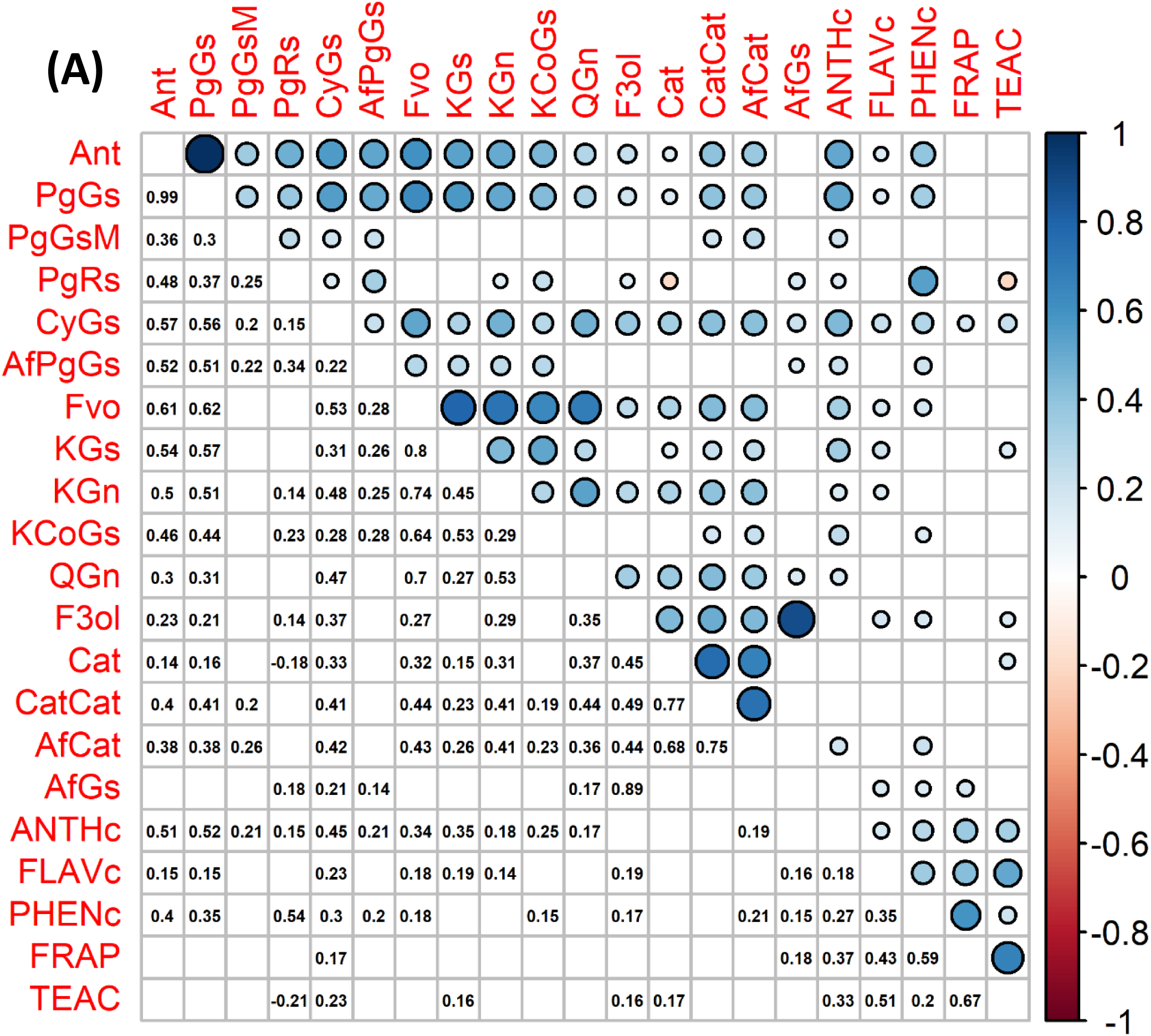
Phenotypic correlations for the traits measured in 2010. (A) Pearson phenotypic correlations (P < 0.05). Correlation values ‘r’ are represented with colour circle in upper right triangle. Scale combines circle size (small circle: correlation near 0; large circle: correlation near 1) and colour gradient from red (negative correlation) to blue (positive correlation). Pearson correlation values are indicated in lower left triangle. Only significant correlations are represented (P < 0.05). A white box represents a non-significant correlation. Diagonal values are not represented. Ant, total anthocyanins; PgGs, Pelargonidin-3-glucoside; PgGsM, Pelargonidin-3-glucoside-malonate; PgRs, Pelargonidin-3-rutinoside; CyGs, Cyanidin-3-glucoside; AfPgGs, (epi)Afzelechin-pelargonidin-3-glucoside; Fvo, total flavonols; KGs, Kaempferol-glucoside; KGn, Kaempferol-glucuronide; KCoGs, Kaempferol-coumaryl-glucoside; QGn, Quercetin-glucuronide; F3ol, total flavan-3-ols; Cat, Catechin; CatCat, (epi)Catechin dimers; AfCat, (epi)Afzelechin-(epi)catechin dimers; AfGs, (epi)Afzelechin-glucoside; ANTHc, anthocyanins (colorimetry); FLAVc, flavonoids (colorimetry); PHENc, phenolics (colorimetry); FRAP, antioxidant (FRAP assay); TEAC, antioxidant (TEAC assay).

Correlations between phenotypic values obtained by colorimetric assays commonly used for measuring anthocyanins (ANTHc), flavonoids (FLAVc), phenolics (PHENc) and antioxidants (FRAP and TEAC) and values obtained for individual flavonoid compounds measured by LC-ESI-MS were also calculated for 2010 (Figure 1A) and for 2011 (Supplemental Figure S1A). For 2010, ANTHc showed indeed high correlations with PgGs (r=0.52) and CyGs (r=0.45), but weak correlations (r=0.15 to 0.35) with other anthocyanins and weak or no significant correlations with flavonols or flavan-3-ols. Strikingly, FLAVc was poorly correlated with all the flavonoid compounds (r=0.14 to 0.23) while PHENc showed a high correlation (r=0.54) only with PgRs and with FRAP and TEAC (r=0.43 and 0.51, respectively). FRAP and TEAC values were highly correlated (r=0.67) but more weakly correlated with all the other traits except PHENc and FLAVc. Similar results were obtained in 2011 for all the traits, except for the eventual odd results such as the high correlation between PHENc and PgRs which was not reproduced. Interestingly, while correlations between compounds and traits were almost exclusively positive in 2010, weak but significant negative correlations could be observed in 2011, for example between the flavan-3-ol AfGs and some anthocyanins (PgGsM, PgRs) and between ANTHc and some flavonols (KCoGs, QGn) and flavan-3-ols (Cat, CatCat).

We further represented the most significant Pearson correlations between the traits (r > 0.3) as a phenotypic correlation network for 2010 (Figure 1B) and for 2011 (Supplemental Figure S1B). The correlation network highlighted the strong relationships between the compounds in each chemical class and between anthocyanins and flavonols. It also pinpointed the weak correlations between colorimetric and metabolic traits.

**Figure 1B.**
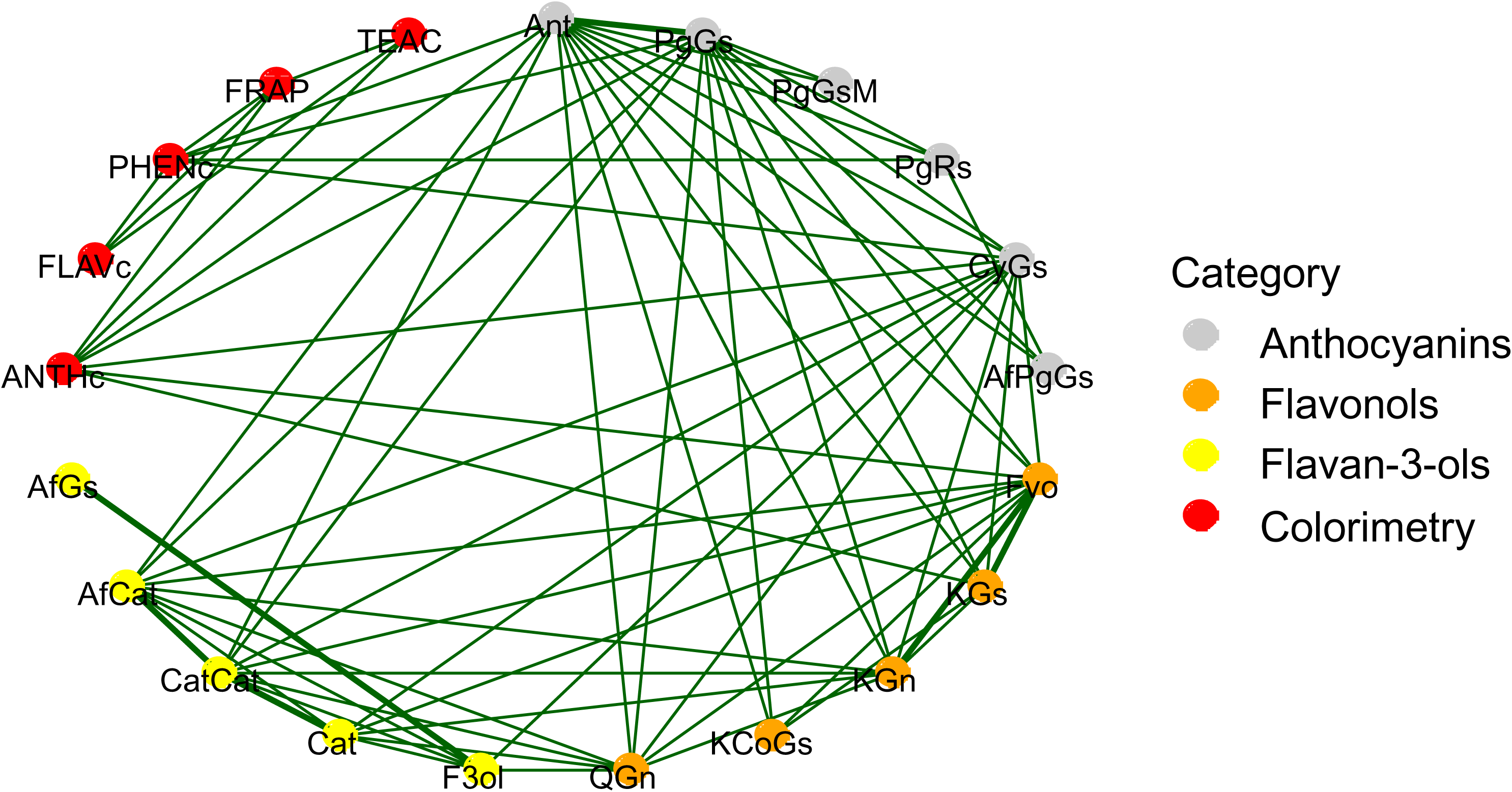
Phenotypic correlations for the traits measured in 2010. (B) Pearson phenotypic correlation network (P < 0.05). Traits are represented as nodes and coloured according to chemical families [Anthocyanins (grey), Flavonols (orange), Flavan-3-ols (yellow)] or to colorimetric assays (red). Positive (green) and negative (red) correlations with absolute values r > |0.3| are represented as links between nodes. The thickness of the links depends on the correlation values; the more the correlation value is high, the more the link is thick. Only significant correlations are represented (P < 0.05).

Scatter plot analysis of trait phenotypic values in 2010 and 2011 (Figure 2) further revealed that ANTHc values had the highest correlation between the two years (r=0.72). High correlations were also observed for the anthocyanin PgGs (r=0.45), the flavonols KGs (r=0.49) and KGn (r=0.42), the flavan-3-ol AfGs (r=0.74) and the ANTHc (r=0.52). Noteworthy, poor correlations (r<0.1) were observed for compounds such as CyGs, QGn and AfCat, underlining the high incidence of environmental conditions on flavonoid accumulation in strawberry fruit.

**Figure 2.**
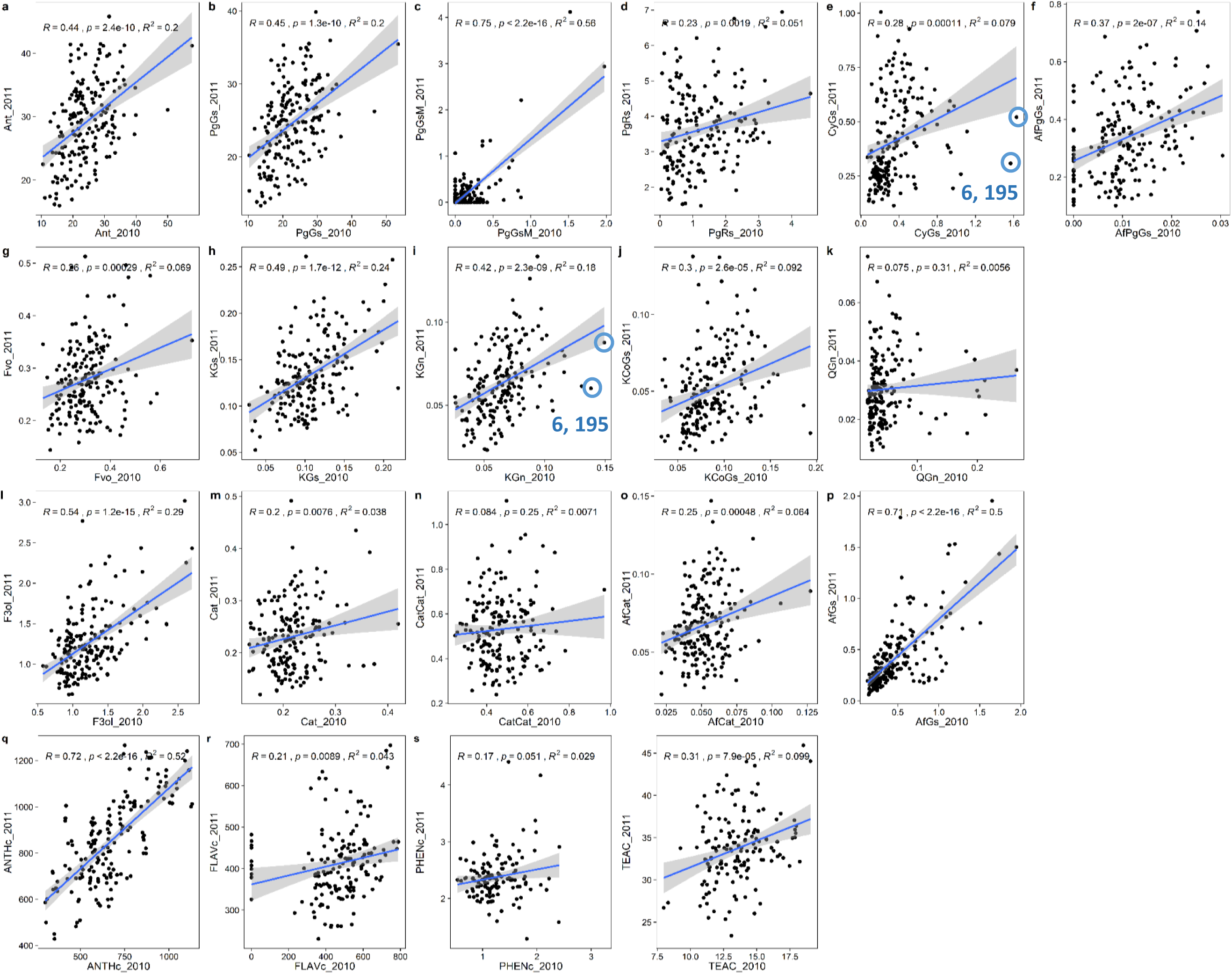
Correlations between the two years. Scatter plots for all traits measured on the two years. The blue line represents the linear regression with associated 95% confident interval. Correlation ratio (R), percentage of variance (R²) and p (p) value of the Pearson test. When p < 0.05, metabolites contents between the two years are correlated. Blue circles indicate examples of genotypes showing extreme values in 2010. Ant, total anthocyanins; PgGs, Pelargonidin-3-glucoside; PgGsM, Pelargonidin-3-glucoside-malonate; PgRs, Pelargonidin-3-rutinoside; CyGs, Cyanidin-3-glucoside; AfPgGs, (epi)Afzelechin-pelargonidin-3-glucoside; Fvo, total flavonols; KGs, Kaempferol-glucoside; KGn, Kaempferol-glucuronide; KCoGs, Kaempferol-coumaryl-glucoside; QGn, Quercetin-glucuronide; F3ol, total flavan-3-ols; Cat, Catechin; CatCat, (epi)Catechin dimers; AfCat, (epi)Afzelechin-(epi)catechin dimers; AfGs, (epi)Afzelechin-glucoside; ANTHc, anthocyanins (colorimetry); FLAVc, flavonoids (colorimetry); PHENc, phenolics (colorimetry); TEAC, antioxidant (TEAC assay).

### QTL analysis

We used the same linkage maps as those previously described in Lerceteau-Köhler et al. (20) with the additional markers described in Ring et al. (19). The markers covered the 28 expected linkage groups (LG) for the female (f) and male (m) linkage maps with two linkage groups (IV-d-f and III-c-m) represented each by two groups (IV-d1-f and IV-d2-f, III-c1-m and III-c2-m) and with five linkage groups anchored (IV-X1 and IV-X2) or not (F30, M41, M44) to one of the seven homoeology groups (HG). To decipher the genetic architecture of the flavonoid compounds, we performed QTL analyses on each of the three replicates of the two successive years (2010 and 2011). Colorimetric traits were analyzed for each year.

Linkage analysis on female and male maps revealed a total of 152 mQTLs for the flavonoid compounds, all of which were identified for the first time from cultivated strawberry, and of 26 QTLs for the antioxidant- and colour-related traits measured by colorimetry (Table 2 and Figure 3). Similar number of mQTLs was identified for each chemical class of the 13 flavonoid compounds analyzed. The anthocyanins (5 compounds) displayed the highest number of mQTLs (57 mQTLs) followed by the flavan-3-ols (4 compounds; 53 mQTLs) and by the flavonols (4 compounds; 42 mQTLs). Remarkably, the number of mQTLs and QTLs mapped on the female and male maps was almost identical for the anthocyanins (28 for the female *vs.* 29 for the male) and for the flavan-3-ols (26 for the female *vs.* 27 for the male) but was uneven for the flavonols (25 for the female *vs.* 17 for the male) and for the colorimetric traits (11 for the female *vs.* 15 for the male).

**Figure 3A.**
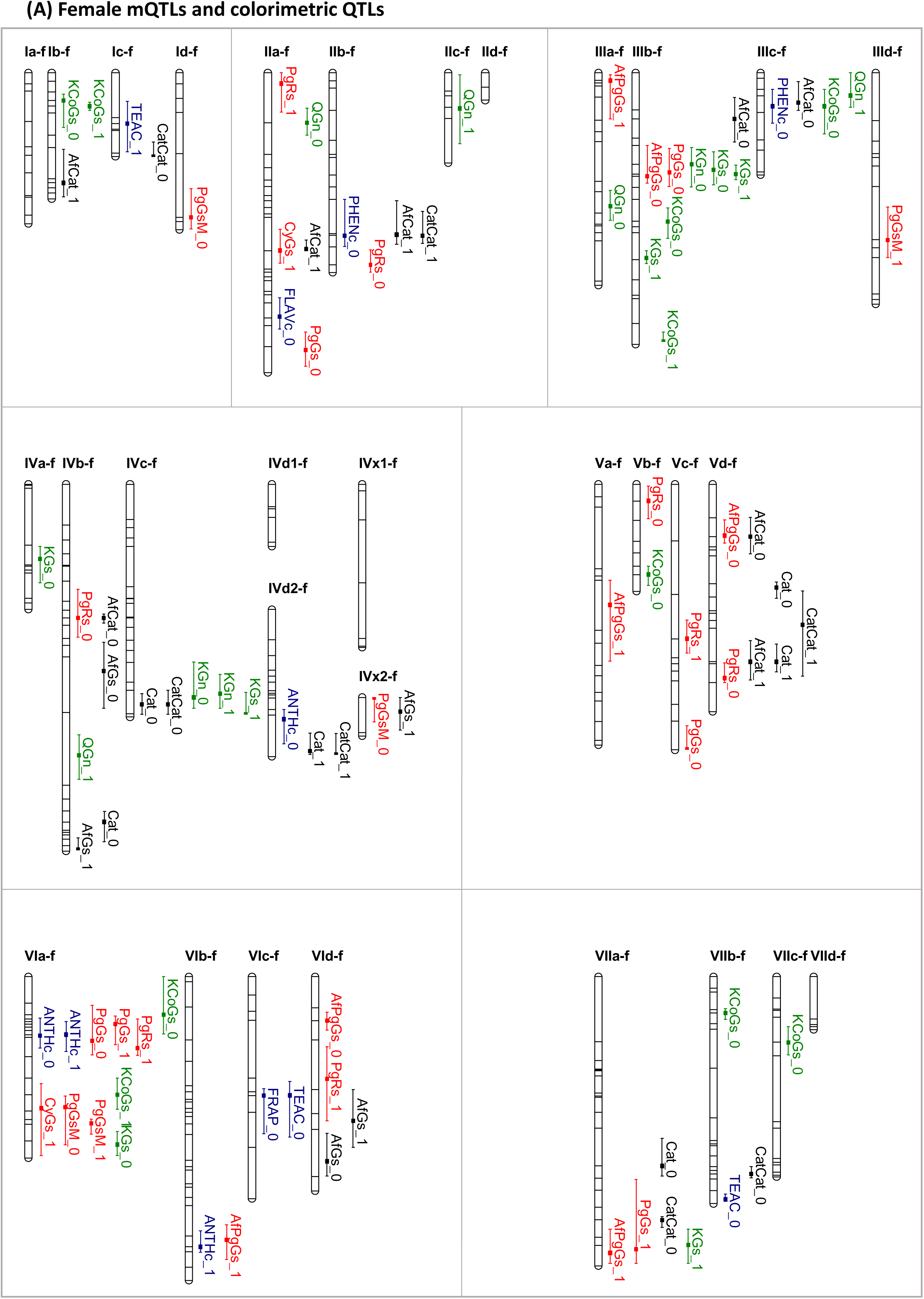
Distribution of QTLs detected in ‘Capitola’ (female). QTLs were detected by QTL Cartographer and are represented by bars of different colours. The different classes of flavonoids and the colour- and antioxidant-related traits are each represented by a single colour Names of mQTLs and colorimetric QTLs are detailed in Supplemental Table S2.

**Figure 3B.**
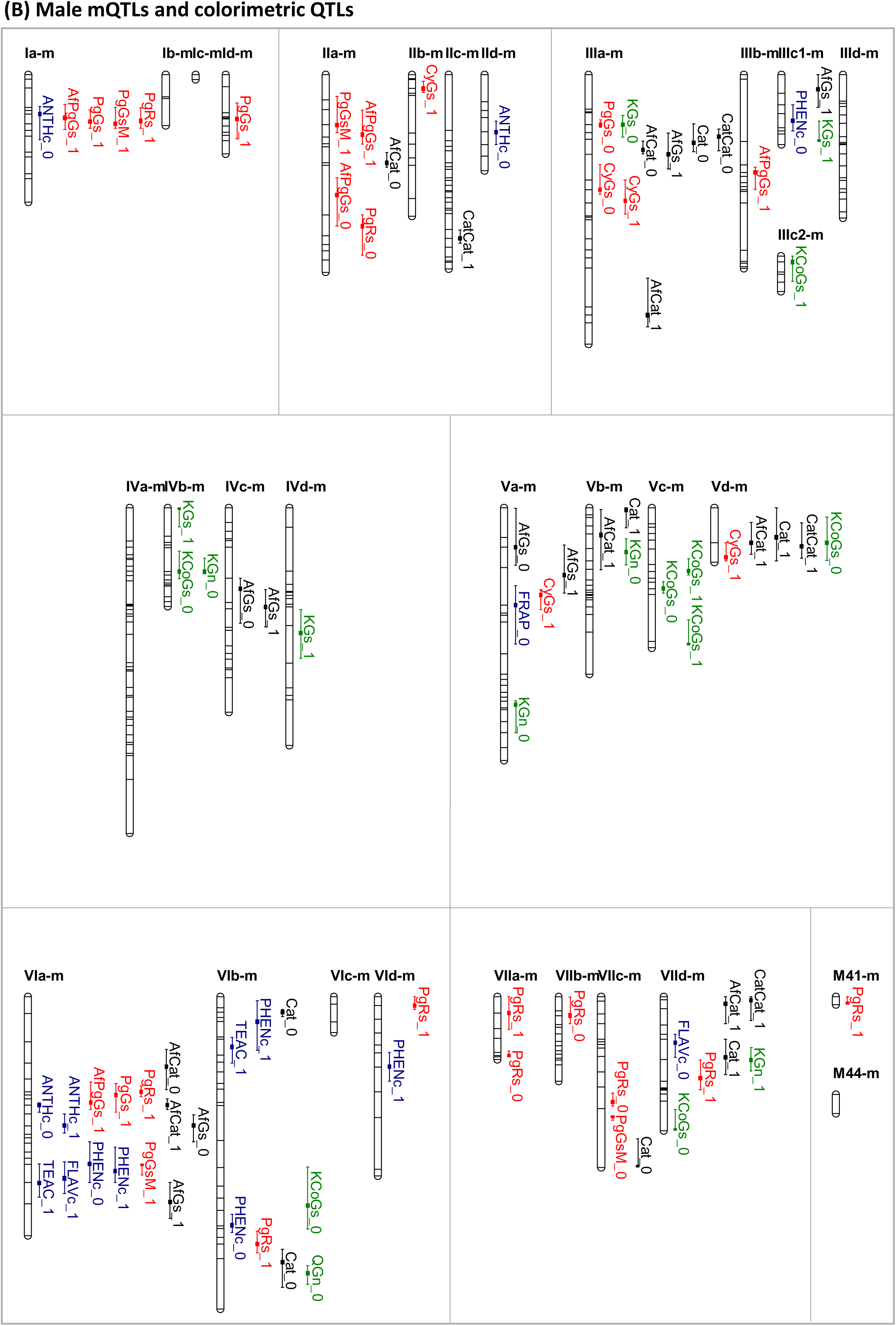
Distribution of QTLs detected in ‘CF1116’ (male). QTLs were detected by QTL Cartographer and are represented by bars of different colours. The different classes of flavonoids and the colour- and antioxidant-related traits are each represented by a single colour. Names of mQTLs and colorimetric QTLs are detailed in Supplemental Table S3.

**Table 2.**
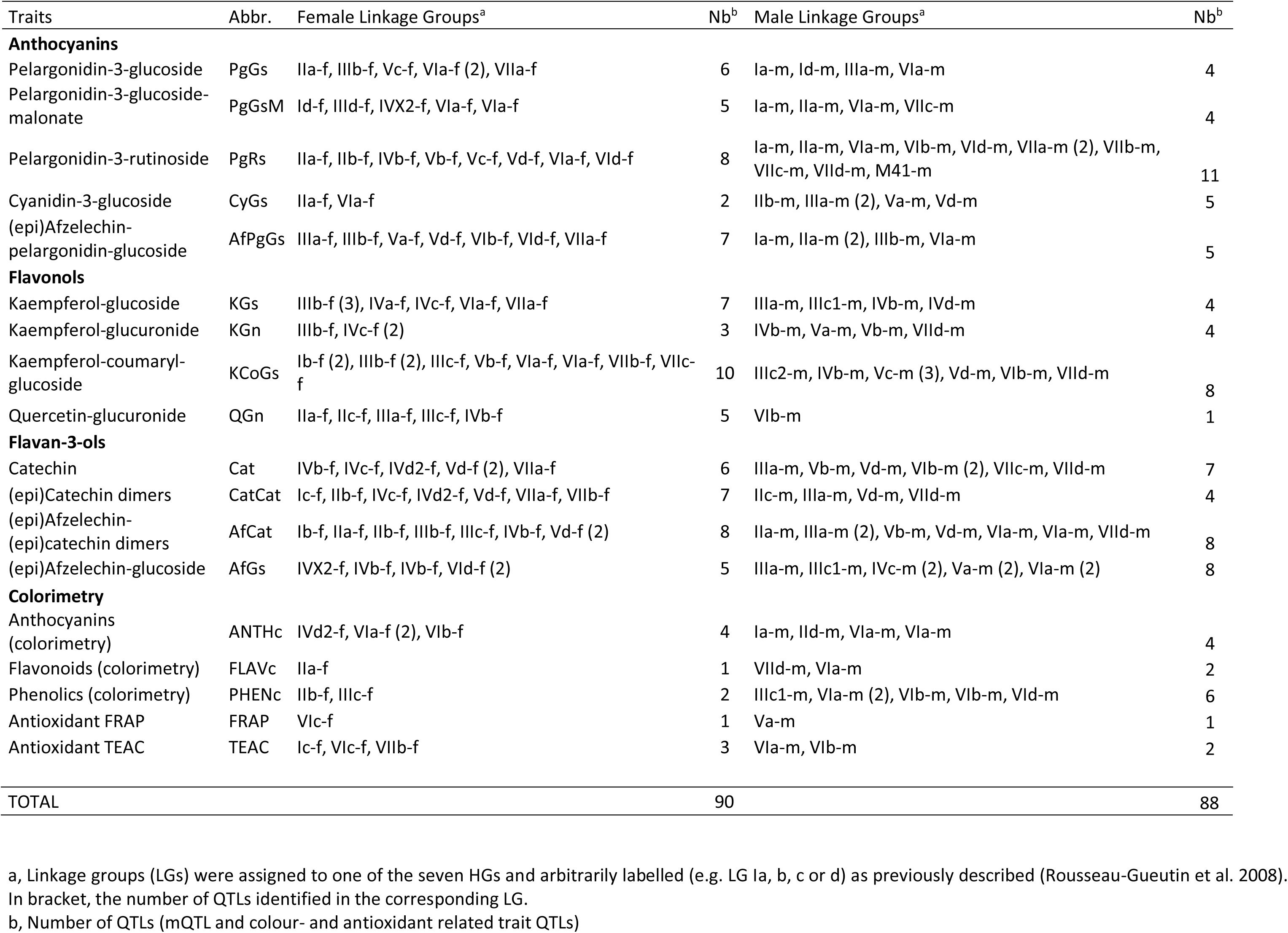
Location of significant QTLs detected for all traits based on CIM analysis with LOD > LOD threshold (significance level α = 0.10).

Because cultivated strawberry is an octoploid species, each of the seven homoeology groups (HGs) has four linkage groups (LGs). LGs were assigned to one of the seven HGs and arbitrarily labelled (e.g. LG Ia, b, c or d) as previously described (35). On both female and male maps, mQTLs were identified for all the HGs (Table 3, Figure 3 and Supplemental Tables S2, S3, S4). Interestingly, the colorimetric QTLs were mostly localized on a unique HG (HG VI) in which 4 QTLs (female) and 10 QTLs (male) were detected whereas none, one or at most two QTLs were mapped on the other HGs (Figure 3). Moreover, within a given HG, mQTLs and colorimetric QTLs were unequally distributed on the various LGs. For example, on the female map, 9 mQTLs among which 6 mQTLs for flavonols (kaempferol derivatives) were detected on LG IIIb while only two were detected on LG IIIa and one on LG IIId. At the opposite, on the male map, only one mQTL was detected on LG IIIb while 9 mQTLs, among which 5 mQTLs for flavan-3-ols, were detected on LG IIIa. A striking result was that in both female and male maps, the largest number of flavonoid mQTLs and colour- and antioxidant-related QTLs (11 on female and 14 on male maps) was detected on the LG VIa (Figure 3) making this linkage group a likely target for the improvement of nutritional quality of strawberry fruit.

**Table 3.**
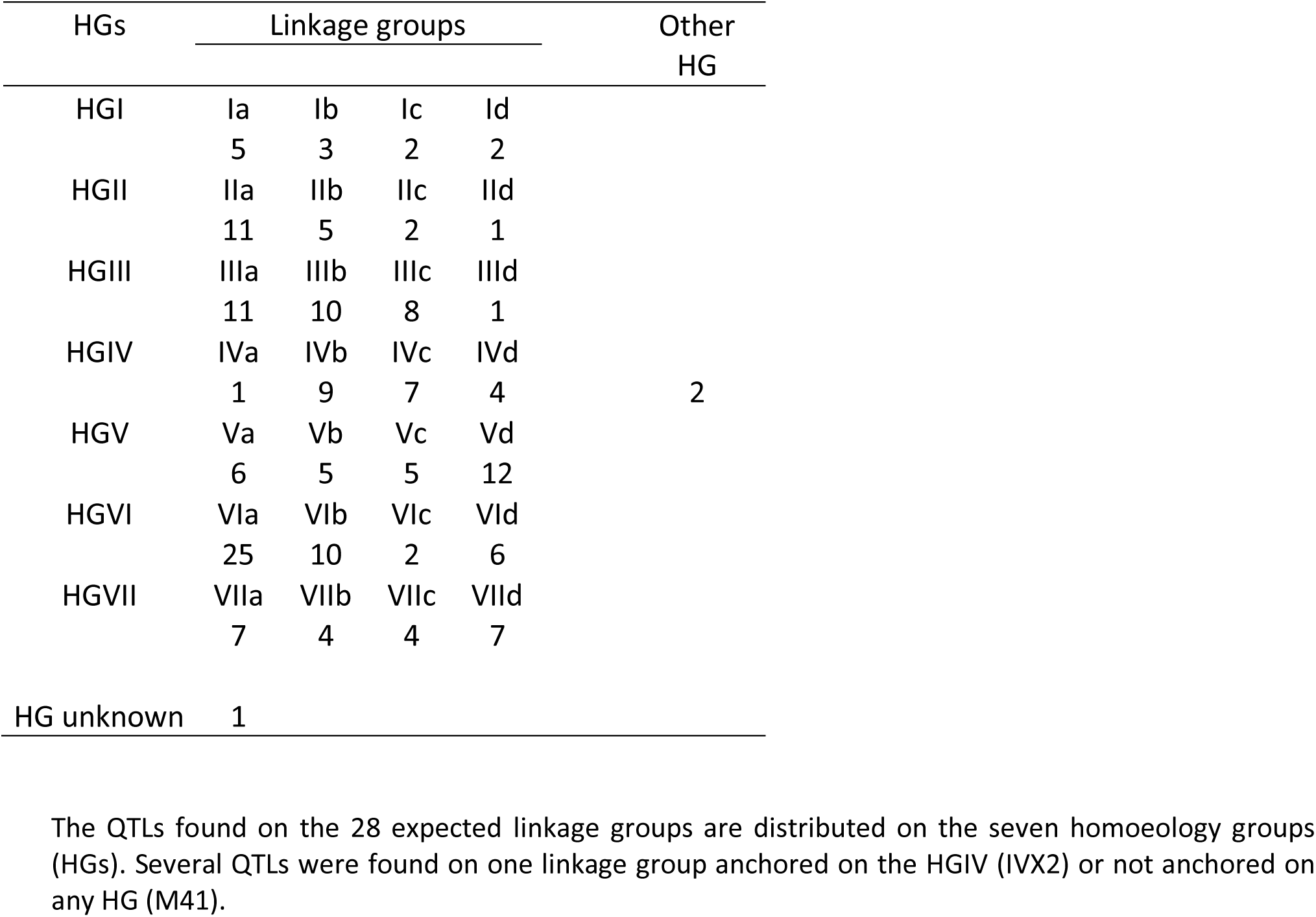
Distribution of QTLs according to the homoeology groups (HG) and linkage groups.

Interestingly, the analysis of the distribution and effects of the mQTLs (Table 2 and Supplemental Tables S2, S3, S4) highlighted large differences between the compounds belonging to a same chemical class. For the most abundant anthocyanin compound, which is pelargonidin-3-glucoside, 6 mQTLs were detected on female map and 4 on male map. Among them, one major PgGs mQTL localized on LG IIIa from male map (LOD values of 3.7-7.4) could explain 14 to 27% of the PgGs variance (R²) in 2010 (Supplemental Tables S2, S3). For the second most abundant anthocyanin compound, pelargonidin-3-rutinoside, which displayed considerable variations in the progeny (Table 1 and Supplemental Table S1), large number of mQTLs was detected on both female (8 mQTLs) and male (11 mQTLs) maps. Though the effect of most PgRs mQTLs on the PgRs phenotypic variation was low, one major mQTL detected on LG VIIb from the male map (LOD value of 4.4) explained 36% of the variance (R²) in 2010. In addition, this PgRs mQTL was co-localized with a major pelargonidin-3-glucoside malonate (PgGsM) (LOD value of 4.5) that also explained 36% of the phenotypic variation (R²) in 2010. Noteworthy, several PgRs mQTLs were localized on each of the four different linkage groups from HG VII on the male map (Figure 3), thus highlighting the possible contribution of each *F.* x *ananassa* subgenome from HG VII to the control of PgRs content. The analysis of the co-localization of anthocyanin mQTLs additionally pinpointed several genomic regions to be considered when breeding for strawberry fruit colour improvement. On the male map, mQTLs for four anthocyanin compounds derived from pelargonidin (PgGs, PgRs, PgGsM, AfPgGs) were co-localized on LG Ia together with the total anthocyanin content measured by colorimetric assay (ANTHc) (Figure 3). Likewise, on both the female and the male maps, different clusters of anthocyanin-related mQTLs were identified on LG VIa (Figure 3). Among them was a cluster of mQTLs for PgGs and PgRs (female map) or for PgGs, PgRs and AfPgGs (male map) that co-localized with ANTHc QTLs and a cluster of overlapping mQTLs for PgGsM and CyGs (female map).

Regarding the flavonols, the most complex genetic architecture was that of the kaempferol-coumaryl-glucoside (KCoGs), with 10 mQTLs on the female map and 8 mQTLs on the male map, followed by the kaempferol-glucoside (KGs), with 7 mQTLs on the female map and 4 mQTLs on the male map (Table 2). Among them, a remarkable mQTL cluster was identified on female LG IIIb where mQTLs for kaempferol derivatives (KGs, KGn) overlapped with those of the anthocyanin PgGs and AfPgGs (Figure 3). Moreover, the female KGs mQTL on LG IIIb was a major mQTL (LOD value of 4.6-5.7) that accounted for 18 to 23% of the explained variance (R²). Major mQTLs were also identified in 2010 for the quercetin-glucuronide (QGn) on the female map on LG IIa (LOD value of 4.5; R² of 20%) and on LG IVb (LOD value of 3.9; R² of 20%) and on male map on LG VIb (LOD value of 4.8; as much as 44% of the variance explained).

As for the flavan-3-ols, the most numerous mQTLs were detected for epiafzelechin derivatives, including 8 female and 8 male mQTLs for (epi)Afzelechin-(epi)catechin (AfCat) dimers and 5 female and 8 male mQTLs for (epi)Afzelechin-glucoside (AfGs). None of the flavan-3-ols mQTLs exhibited large explained phenotypic variance (R² > 20%) in 2010 except the female AfCat mQTL located on LG IIIc (LOD value of 3.6; R² of 22%) and Cat mQTL located on LG Vd (LOD value of 4.8; R² of 22%). The most interesting flavan-3-ols mQTL cluster was that of catechin (Cat, CatCat) and epiafzlechin (AfGs, AfCat) derivatives that overlapped on male map on a narrow genomic region from LG IIIa.

Surprisingly, while 8 ANTHc QTLs and 8 PHENc QTLs were identified, few QTLs were discovered for the fruit antioxidant capacity measured by FRAP (2 in total) and TEAC (5 in total) assays. Noteworthy, the colorimetric QTLs, PHENc, FLAVc and TEAC were clustered on LG VIa on female map (Figure 3).

## Discussion

Small berry fruits, including strawberry, are an importance source of antioxidants in our diet.(4, 36). Antioxidant capacity of strawberry fruit depends not only on antioxidants such as vitamin C (11) but also on its composition in polyphenolics, which contribute to both its sensorial and nutritional quality.(5,6,23). Thanks to the recent advances in the analysis of specialized metabolism (22), it is now possible to breakdown complex traits such as fruit colour and antioxidant capacity into more discrete traits controlling variations in individual chemical compounds. We investigated here the fruit flavonoid and antioxidant content of a pseudo full-sibling F_1_ population obtained from a cross between the variety ‘Capitola’ and the advanced line ‘CF1116’. The two parents, ‘Capitola’ and ‘CF1116’, display many contrasting fruit quality traits.(26). ‘Capitola’ produces large fruits with low sugar and high acidity while ‘CF1116’ produces small fruits with high sugar and lower acidity. In addition, the segregating population issued from ‘Capitola’ and ‘CF1116’ displays considerable variation in fruit colour, as shown by the differences in the Lab colour space values and total anthocyanin content.(20). In a previous work, we mapped quantitative trait loci (QTLs) for various fruit quality traits related to fruit development, texture, sugar and organic acid contents as well as fruit colour.(20). Mapping flavonoid metabolic QTLs (mQTLs) to specific linkage groups in the octoploid cultivated strawberry (21), as done in this study, and comparative analysis of their localization with that of other fruit quality QTLs previously described (20) will facilitate the identification of genomic regions that can be targeted through marker-assisted selection (MAS) for breeding superior strawberry varieties with enhanced sensorial and nutritional quality.

### The ‘Capitola’ x ‘CF1116’ population displays a large variability in flavonoid composition

Anthocyanins are water-soluble flavonoid pigments that constitute the major flavonoid compounds found in strawberry fruit. They are responsible for the red colour of strawberry fruit and strongly contribute to its antioxidant capacity.(23). In the parents and progeny studied, we found that the major anthocyanin compound in the fruit is pelargonidin-3-glucoside (∼84-90 % of the anthocyanins) followed by pelargonidin-3-rutinoside while minor anthocyanin compounds (cyanidin-3-glucoside, pelargonidin-3-glucoside-malonate, (epi)Afzelechin-pelargonidin-glucoside) were also detected (Table 1 and Supplemental Talbe S1). While anthocyanins were by far the major contributors (∼95-97 %) to the flavonoids detected in the ‘Capitola’ and ‘CF1116’ parents and progeny, other flavonoids belonging to the flavan-3-ols (∼2 to 4 %) and flavonols chemical classes were also detected and quantified (Table 1 and Supplemental Table S1). Although flavonols represent a very minor fraction (1 %) of the flavonoids in cultivated strawberry (23), mainly in the form of kaempferol and quercetin derivatives (Table 1 and Supplemental Table S1), their intake has a health-promoting effect and reduces the risk of several diseases.(5, 7).

The relative contributions of individual compounds to their respective chemical classes are in agreement with previous findings for cultivated strawberry.(12,19,23). However, they may contrast with other results published for the wild diploid woodland strawberry *F. vesca*.(14). In this species, for example, the major anthocyanin compounds are both pelargonidin-3-glucoside and cyanidin-3-glucoside, which may respectively contribute to 50% and 40% of the fruit anthocyanins.(12, 14). In our segregating population, most of the variations among the most extreme genotypes in the progeny were in the 4 to 10-fold range, an example of which is the ∼5-fold variation in pelargonidin-3-glucoside content and the ∼4-fold variation in flavonols; it is worth mentioning that values for these compounds were similar in the parents regardless of the large variations in the progeny. Much higher variations were observed in the progeny for the anthocyanins cyanidin-3-glucoside and pelargonidin-3-rutinoside (23-fold and 114-fold, respectively) and for the flavan-3-ol (epi)afzelechin-glucoside (17-fold). Such variations in individual flavonoids may affect the nutritional value of the fruit because of the synergetic effect of anthocyanins and flavonols on health.(37). They may also affect fruit sensorial quality because anthocyanin composition will likely affect fruit colour intensity and hue.(38, 39).

### High connectivity between individual flavonoid compounds within each chemical class but low correlation with antioxidant traits measured by colorimetric assays

Not surprisingly given that they share common biosynthetic pathways, the various anthocyanins were positively correlated with each other, as were the flavonol and the flavan-3-ol compounds (Figure 1A and B and Supplemental Figure S1). In addition, high positive correlation values were observed between anthocyanins and flavonols, highlighting the tight connection between these chemical flavonoid classes, which share naringenin and dihydrokaempferol as common precursors. Co-regulations of different enzymes involved at various steps of the flavonoid pathway may additionally explain the parallel variations of the various flavonoids in the progeny studied. Such co-regulations have been well documented in various plant species, including strawberry.(40, 41).

The correlations between the total flavonoids, phenolics, anthocyanins and antioxidant trait values estimated by colorimetric methods (FLAVc, PHENc, ANTHc, FRAP and TEAC) and the various flavonoids determined by LC-MS-ESI produced some unexpected results. While anthocyanins contents were well correlated with ANTHc, the correlations with FLAVc, FRAP, and TEAC were poor (r < 0.3). The weak correlations of anthocyanins and other flavonoids with FRAP and TEAC can be explained by the differences in the metabolites detected by colorimetric assays and by LC–ESI–MS analysis. In addition, both FRAP and TEAC measure total antioxidants, among which the flavonoids, but also additional phenolics such as the ellagic acid that has high antioxidant and nutritional values.(42). Indeed, pelargonidin-3-glucoside, which is the main flavonoid compound, only accounts for ∼25% of the total strawberry antioxidant capacity.(23). Likewise, PHENc, which is highly correlated with FRAP (Figure 1A and B), is not restricted to flavonoids but also estimates total phenolics.(30). More surprising are the weak correlations between the total flavonoid content measured by LC–ESI–MS and that estimated by FLAVc assay, which questions the use of colorimetric methods commonly used for assessing fruit nutritional quality and the interpretations of their results.

In addition to measuring the variations in individual flavonoid compounds in the ‘Capitola’ x ‘CF1116’ progeny for a given year, we measured for each trait the correlation values between the two successive years (Figure 2). High correlations were observed for several traits, including colour-related traits (ANTHc, PgGs content) but also flavonols (kaempferol derivatives KGs and KGn) and flavan-3-ol (AfGs), highlighting the strong genetic control of flavonoid composition already observed in *F. vesca*.(14). In our experimental conditions where plants were cultivated under plastic tunnel and therefore subjected to natural climatic variations, a likely explanation for the poor correlations observed for some traits, an example of which is the CyGs anthocyanin (Figure 2), is the effect of environmental conditions. Sensitivity of flavonoid metabolism to environmental conditions is well documented.(43). Additionally, because poor correlations were mostly due to few individuals of the progeny showing contrasted and extreme phenotypic values in 2010 (e.g. the individuals no. 6 and 195), a strong genotypic effect in the phenotypic plasticity i.e. the response to environmental variations (here year of study), is likely.(44).

### Flavonoid mQTL mapping and comparison with fruit quality QTLs previously identified

Evaluation and genetic mapping of both specialized (polyphenolics, flavor components) and primary (sugars, organic acids) metabolites are indispensable to identify the genetic architecture of fruit sensorial and nutritional quality in cultivated strawberry.(20,45–47). In addition, genetic studies allow pinpointing genotypes that can be used in a breeding scheme for the genetic improvement of strawberry. In the post-genomic era, to efficiently harness the available strawberry diversity and translate these findings into crop improvement, genetic studies have to be extended to whole populations displaying large phenotypic and genetic diversity.(10), which are typically bi-parental populations in cultivated strawberry.(20,45–47). Using such a bi-parental population issued from a cross between two parents with contrasted sensory trait values (20) and by additionally breaking down the flavonoid composition to its individual chemical components (22), we were able to map mQTLs for each chemical compound from the three flavonoid chemical classes detected in strawberry fruit. The analysis of three replicates over two successive years for each trait allowed us to identify some robust QTLs for specialized metabolism. Additional QTLs that were not significantly detected in the two successive years nevertheless displayed a significant effect from one year to another. As mentioned above for trait correlations between years, such QTLs are likely more sensitive to the environmental conditions, which may considerably vary when plants are grown in natural conditions.

An example of a major mQTL (14 to 27% of the variance explained) that is possibly sensitive to environmental conditions is the mQTL for pelargonidin-3-glucoside, the most abundant anthocyanin compound, which was detected on male linkage group LG IIIa for the three replicates in 2010 but not in 2011. Because this mQTL is also co-localized with robust QTLs for the colour physical parameters L and b (colour space values), which were previously detected over three successive years (20), it can nevertheless be targeted for fruit colour improvement. Additionally, because we used the same linkage mapping approach and set of markers as in our previous study (20), we could identify a LG IIIa-m gata165c marker linked to both PgGs content and ANTHc and L colour-related traits. This marker is also adjacent to the SSR EFMv029 (allele v029205c) marker, which can therefore be used for the marker-assisted selection (MAS) screening of genotypes-of-interest displaying fruits with enhanced colour and sweetness. Thus, by combining fruit flavonoid mQTL with QTLs previously detected for diverse fruit quality traits (20), we could detect a genomic region that can be targeted for both fruit sensorial and nutritional quality.

Another genomic region that appears on both female and male maps as a major linkage group for the control of nutritional traits related to specialized metabolism and of sensorial traits is LG VIa. In our previous study on strawberry fruit sensorial traits, LG VIa (male and/or female) already appeared as an important linkage group for the control of traits related to primary metabolism such as Soluble Solid Content (SSC), sucrose and glucose contents, pH and malic and citric acid contents. In this case as well, markers common to taste and colour-related traits could be identified. These include the SSR EMF006 (allele v006205c) marker, which is close to the AFLP markers tgta383 and gata170 in male, and the SSR Fvi020 (allele g020175) marker, which is close to the AFLP ccta303 marker in female. Both markers are linked to several taste- and colour-related QTLs (SSC, sucrose, a and b color space values, ANTH) previously identified (20) and to the colour-related PgGs mQTL and ANTHc QTL detected in the present study. These markers can be further used for the early selection of genotypes for improved sensorial (sweetness, colour) and nutritional (anthocyanin content) quality.

In addition to its interest for accelerating breeding through MAS, the QTL mapping strategy can also help deciphering which pathway(s) or enzyme(s) are likely involved in the natural variation of a given trait. In the last decades, reverse genetic studies have successfully been used in strawberry and in other fruit species to investigate the role of regulatory or structural genes in the control of specific steps of the phenylpropanoid pathway. Such studies often rely upon the silencing or overexpression of a candidate gene whose possible role is inferred from its already known function. Among the many studies published for strawberry are the demonstration of the crucial role of MYB transcription factors and associated MYB-bHLH-WD40 complex in the regulation of anthocyanin biosynthesis (48), of the anthocyanidin glucosyltransferase in flavonoid biosynthesis (16), and of the anthocyanidin reductase (ANR) enzyme in the trade-off between anthocyanidin and flavan-3-ols.(17). Precise QTL mapping now gives access to candidate genes underlying natural quantitative variation in a given metabolite, an example of which is the discovery of the role of a peroxidase in the trade-off between lignin and anthocyanin biosynthesis.(19). The analysis of the clustering of mQTLs for compounds derived from the same pathway, e.g. the clusters of pelargonidin derived compounds on female LG VIa, or from different pathways, e.g. the clusters of pelargonidin (anthocyanins) and kaempferol (flavonols) derived products on female LG IIIb, additionally provide cues on which step of the flavonoid pathway is worth investigating. The recent sequencing of the octoploid cultivated strawberry (21) will now give access to the tools necessary to identify the underlying candidate genes and the genetic variations responsible for the disparities in flavonoid-related traits in strawberry.

## Acknowledgements

Thanks for their help to Karine Tallès and Gabriel Jousseaume for fruit harvests and colorimetric measurements.

## Author contribution statement

BD conceived and designed the experiments. AuP conducted hands-on experiments and data collection. AuP, AlP and AG participated in the data collection. LR, TH and WS generated LC-LS data. ML, GV, CR and BD conducted data analysis and performed statistical analysis. CR wrote the original draft. All authors read and approved the final manuscript.

## SUPPORTING INFORMATION

**Supplemental Figure S1.**
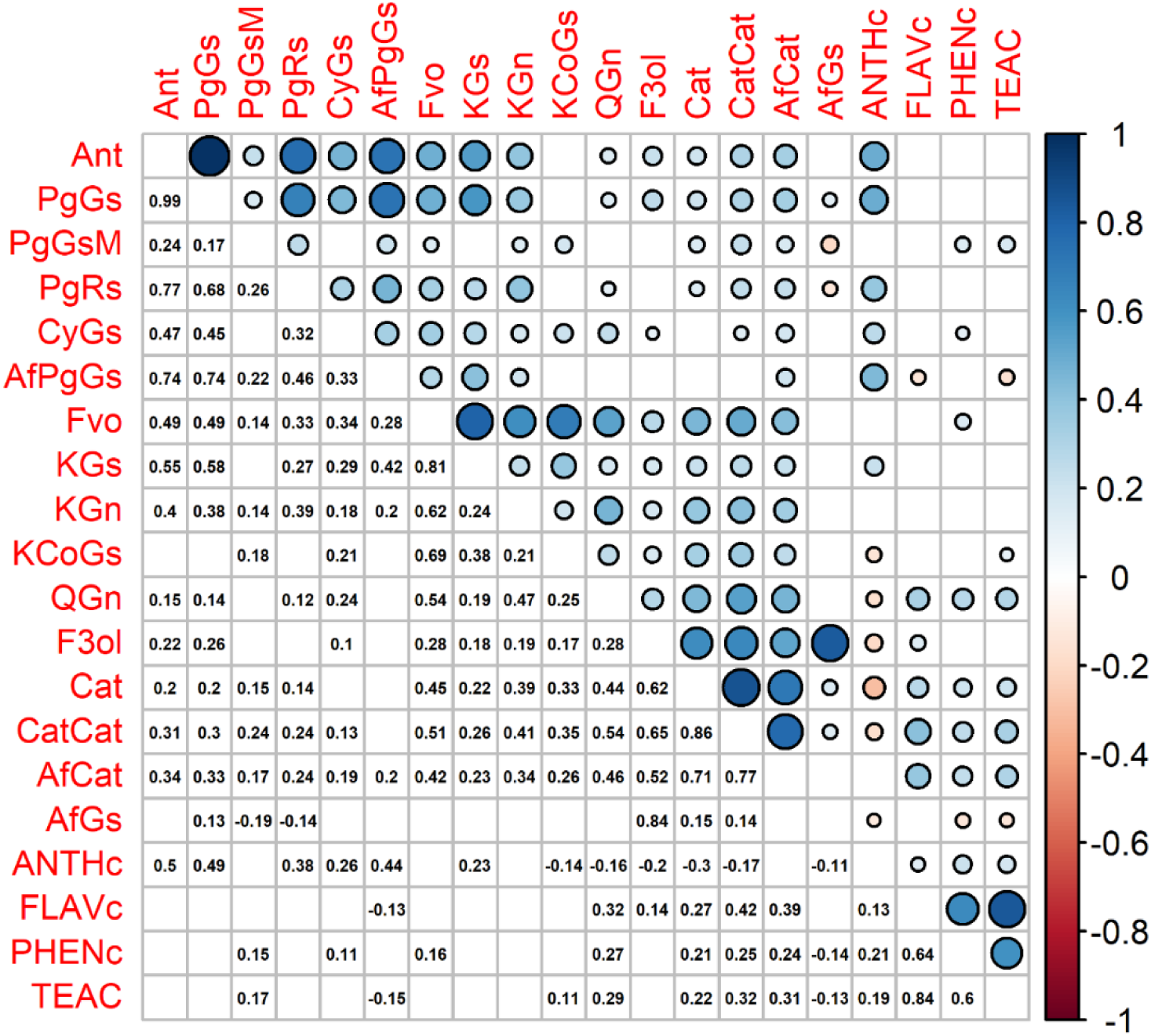
Phenotypic correlations for the traits measured in 2011. **A) Pearson phenotypic correlations (P < 0.05).** Correlation values ‘r’ are represented with colour circle in upper right triangle. Scale combines circle size (small circle: correlation near 0; large circle: correlation near 1) and colour gradient from red (negative correlation) to blue (positive correlation). Pearson correlation values are indicated in lower left triangle. Only significant correlations are represented (P < 0.05). A white box represents a non-significant correlation. Diagonal values are not represented. Ant, total anthocyanins; PgGs, Pelargonidin-3-glucoside; PgGsM, Pelargonidin-3-glucoside-malonate; PgRs, Pelargonidin-3-rutinoside; CyGs, Cyanidin-3-glucoside; AfPgGs, (epi)Afzelechin-pelargonidin-3-glucoside; Fvo, total flavonols; KGs, Kaempferol-glucoside; KGn, Kaempferol-glucuronide; KCoGs, Kaempferol-coumaryl-glucoside; QGn, Quercetin-glucuronide; F3ol, total flavan-3-ols; Cat, Catechin; CatCat, (epi)Catechin dimers; AfCat, (epi)Afzelechin-(epi)catechin dimers; AfGs, (epi)Afzelechin-glucoside; ANTHc, anthocyanins (colorimetry); FLAVc, flavonoids (colorimetry); PHENc, phenolics (colorimetry); FRAP, antioxidant (FRAP assay); TEAC, antioxidant (TEAC assay).

**Figure S1.**
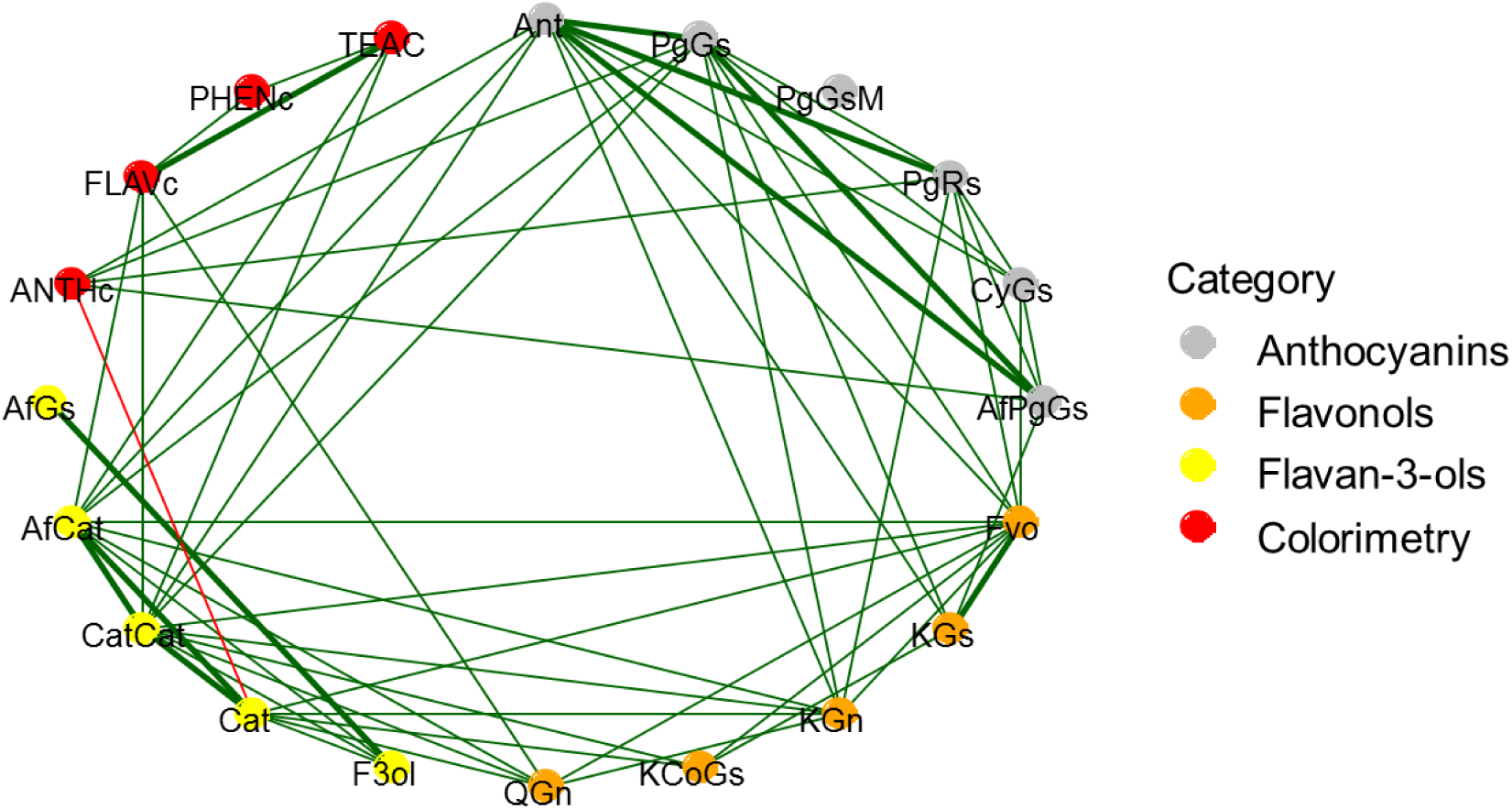
Phenotypic correlations for the traits measured in 2011. **B) Pearson phenotypic correlation network (P < 0.05).** Traits are represented as nodes and coloured according to chemical families [Anthocyanins (grey), Flavonols (orange), Flavan-3-ols (yellow)] or to colorimetric assays (red). Positive (green) and negative (red) correlations with absolute values r > |0.3| are represented as links between nodes. The thickness of the links depends on the correlation values; the more the correlation value is high, the more the link is thick. Only significant correlations are represented (P < 0.05). Ant, total anthocyanins; PgGs, Pelargonidin-3-glucoside; PgGsM, Pelargonidin-3-glucoside-malonate; PgRs, Pelargonidin-3-rutinoside; CyGs, Cyanidin-3-glucoside; AfPgGs, (epi)Afzelechin-pelargonidin-3-glucoside; Fvo, total flavonols; KGs, Kaempferol-glucoside; KGn, Kaempferol-glucuronide; KCoGs, Kaempferol-coumaryl-glucoside; QGn, Quercetin-glucuronide; F3ol, total flavan-3-ols; Cat, Catechin; CatCat, (epi)Catechin dimers; AfCat, (epi)Afzelechin-(epi)catechin dimers; AfGs, (epi)Afzelechin-glucoside; ANTHc, anthocyanins (colorimetry); FLAVc, flavonoids (colorimetry); PHENc, phenolics (colorimetry); FRAP, antioxidant (FRAP assay); TEAC, antioxidant (TEAC assay).

**Supplemental Table 1.** Trait values for ‘Capitola’ and ‘Capitola’ x ‘CF1116’ progeny in 2011.

| Traits | Abbr. | Capitola |  |  | CF1116 |  |  |  | Progeny |  |  | h<sup>2</sup> | Tr |
| --- | --- | --- | --- | --- | --- | --- | --- | --- | --- | --- | --- | --- | --- |
|  |  | mean | s.d. | range | mean | s.d. | range | Cap vs CF | mean | s.d. | range |  |  |
| Anthocyanins |  |  |  |  |  |  |  |  |  |  |  |  |  |
| Total anthocyanins | Ant | 38.52 | 8.37 | 31.65-47.84 | NA | NA | NA | NA | 29.34 | 7.23 | 13.23-53.99 | 0.55 | NA |
| Pelargonidin-3-glucoside | PgGs | 32.26 | 6.91 | 26.65-39.98 | NA | NA | NA | NA | 24.87 | 6.06 | 11.19-45.31 | 0.54 | NA |
| Pelargonidin-3-glucoside-malonate | PgGsM | 0.23 | 0.06 | 0.19-0.30 | NA | NA | NA | NA | 0.15 | 0.34 | 0.00-4.12 | 0.97 | NA |
| Pelargonidin-3-rutinoside | PgRs | 4.97 | 1.15 | 3.96-6.22 | NA | NA | NA | NA | 3.58 | 1.24 | 1.16-7.62 | 0.8 | NA |
| Cyanidin-3-glucoside | CyGs | 0.65 | 0.18 | 0.52-0.85 | NA | NA | NA | NA | 0.41 | 0.19 | 0.11-1.01 | 0.91 | NA |
| (epi)Afzelechin-pelargonidin-glucoside | AfPgGs | 0.41 | 0.08 | 0.32-0.49 | NA | NA | NA | NA | 0.33 | 0.16 | 0.05-0.85 | 0.9 | NA |
| Flavonols |  |  |  |  |  |  |  |  |  |  |  |  |  |
| Total Flavonols | Fvo | 0.27 | 0.05 | 0.21-0.31 | NA | NA | NA | NA | 0.28 | 0.07 | 0.11-0.53 | 0.65 | NA |
| Kaempferol-glucoside | KGs | 0.11 | 0.02 | 0.09-0.13 | NA | NA | NA | NA | 0.13 | 0.04 | 0.05-0.26 | 0.74 | NA |
| Kaempferol-glucuronide | KGn | 0.08 | 0.01 | 0.06-0.09 | NA | NA | NA | NA | 0.06 | 0.02 | 0.02-0.16 | 0.82 | NA |
| Kaempferol-coumaryl-glucoside | KCoGs | 0.06 | 0.02 | 0.04-0.08 | NA | NA | NA | NA | 0.05 | 0.03 | 0.01-0.15 | 0.74 | NA |
| Quercetin-glucuronide | QGn | 0.02 | 0.01 | 0.02-0.03 | NA | NA | NA | NA | 0.03 | 0.01 | 0.01-0.10 | 0.88 | NA |
| Flavan-3-ols |  |  |  |  |  |  |  |  |  |  |  |  |  |
| Total Flavan-3-ols | F3ol | 0.95 | 0.14 | 0.80-1.09 | NA | NA | NA | NA | 1.32 | 0.42 | 0.52-3.02 | 0.87 | NA |
| Catechin | Cat | 0.20 | 0.03 | 0.17-0.22 | NA | NA | NA | NA | 0.24 | 0.06 | 0.12-0.49 | 0.64 | NA |
| (epi)Catechin dimers | CatCat | 0.47 | 0.08 | 0.39-0.54 | NA | NA | NA | NA | 0.55 | 0.16 | 0.24-1.11 | 0.77 | NA |
| (epi)Afzelechin-(epi)catechin dimers | AfCat | 0.06 | 0.01 | 0.05-0.08 | NA | NA | NA | NA | 0.07 | 0.02 | 0.02-0.15 | 0.81 | NA |
| (epi)Afzelechin-glucoside | AfGs | 0.22 | 0.02 | 0.19-0.24 | NA | NA | NA | NA | 0.47 | 0.32 | 0.06-1.95 | 0.96 | NA |
| Colorimetry |  |  |  |  |  |  |  |  |  |  |  |  |  |
| Anthocyanins (colorimetry) | ANTHc | 870.64 | 14.42 | 854.2-881.4 | NA | NA | NA | NA | 831.1 | 209.0 | 393.6-1456.0 | 1.0 | NA |
| Flavonoids (colorimetry) | FLAVc | 301.51 | 19.60 | 280.9-319.9 | NA | NA | NA | NA | 422.7 | 95.5 | 230.6-858.4 | 1.0 | NA |
| Phenolics (colorimetry) | PHENc | 2.13 | 0.55 | 1.72-2.76 | NA | NA | NA | NA | 2.38 | 0.39 | 1.29-4.70 | 0.89 | NA |
| Antioxidant FRAP | FRAP | NA | NA | NA | NA | NA | NA | NA | NA | NA | NA | NA | NA |
| Antioxidant TEAC | TEAC | 26.83 | 2.53 | 24.94-29.70 | NA | NA | NA | NA | 34.41 | 4.77 | 21.45-49.15 | 0.95 | NA |

For each trait, the mean value, standard deviation (s.d.), range and heritability ( $h^2$ ) are described. ‘CF1116’ and transgressive values were not studied in 2011.
Flavonoid metabolites values are expressed as mg-equ/100 g FW assuming a response factor of 1.
Colour- and antioxidant-related traits: ANTHc results are expressed as mg pelargonidin-3-glucoside equivalents/100 g FW. FLAVc results are expressed as $\mu\text{g}$ catechin equivalents/g FW. PHENc results are expressed as mg gallic acid equivalents/g FW. FRAP and TEAC results are expressed as $\mu\text{M}$ Trolox equivalents/g FW. Values are the means of $n = 3$ replicates per genotype.

**Supplemental Table 2.**
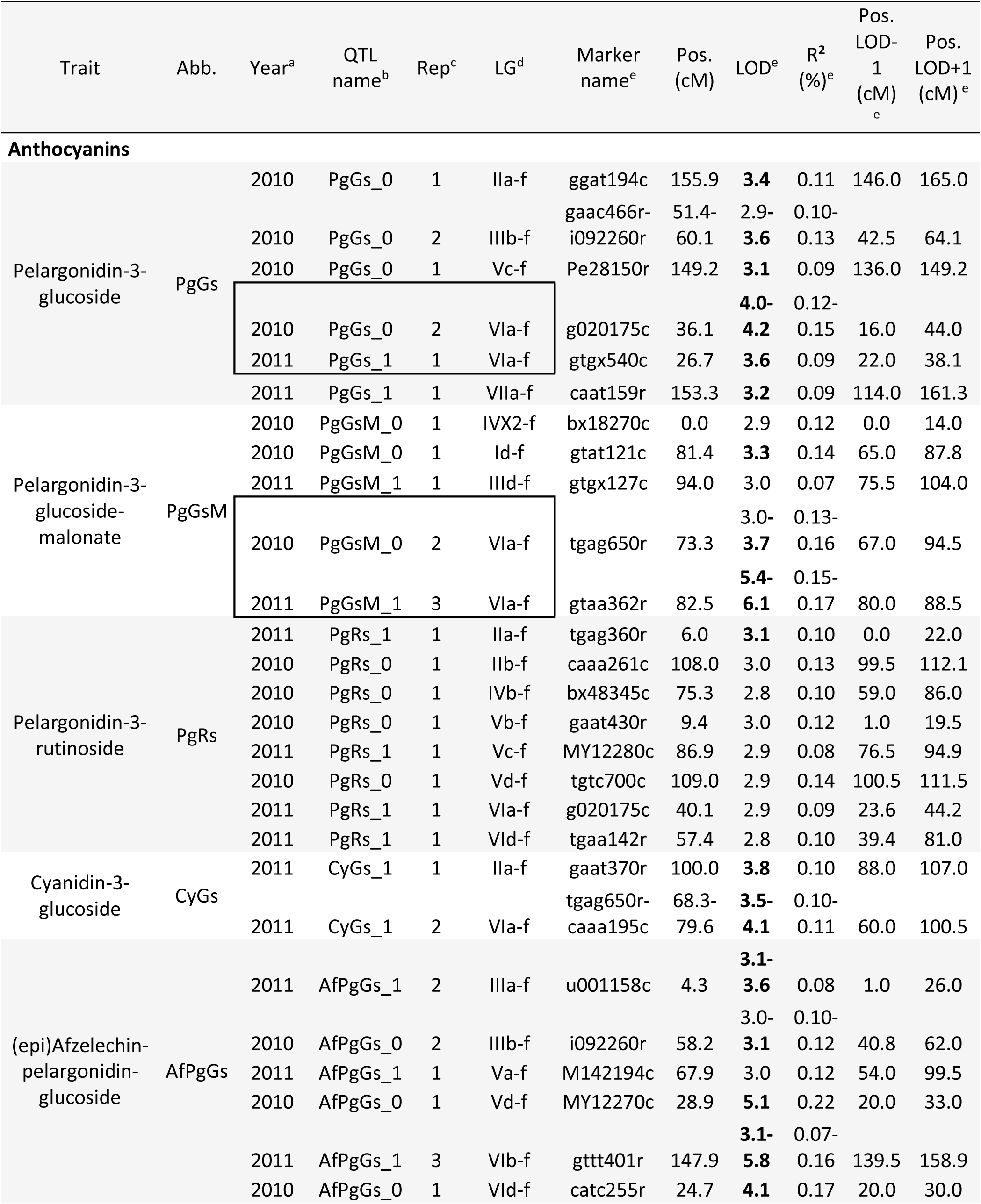
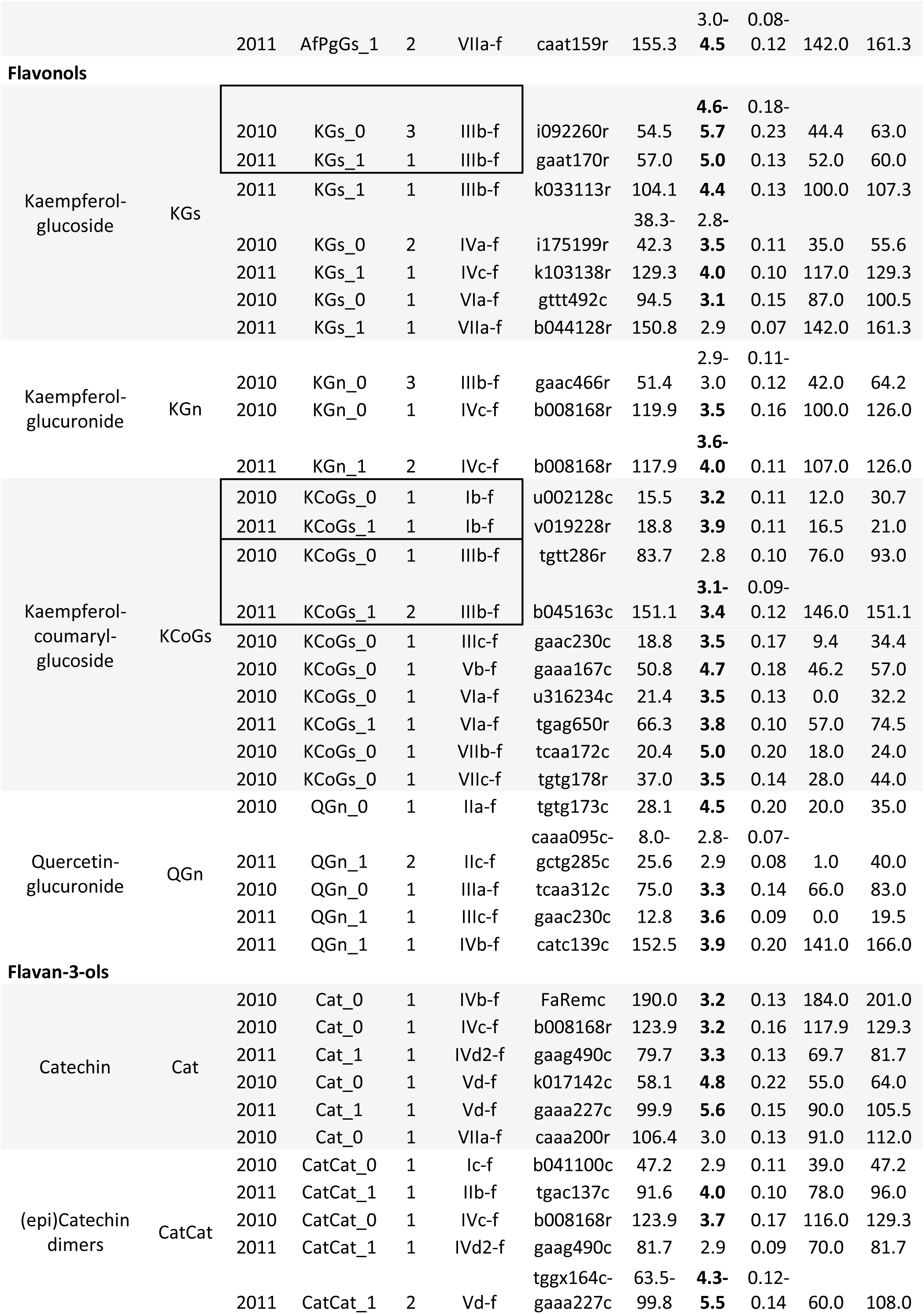
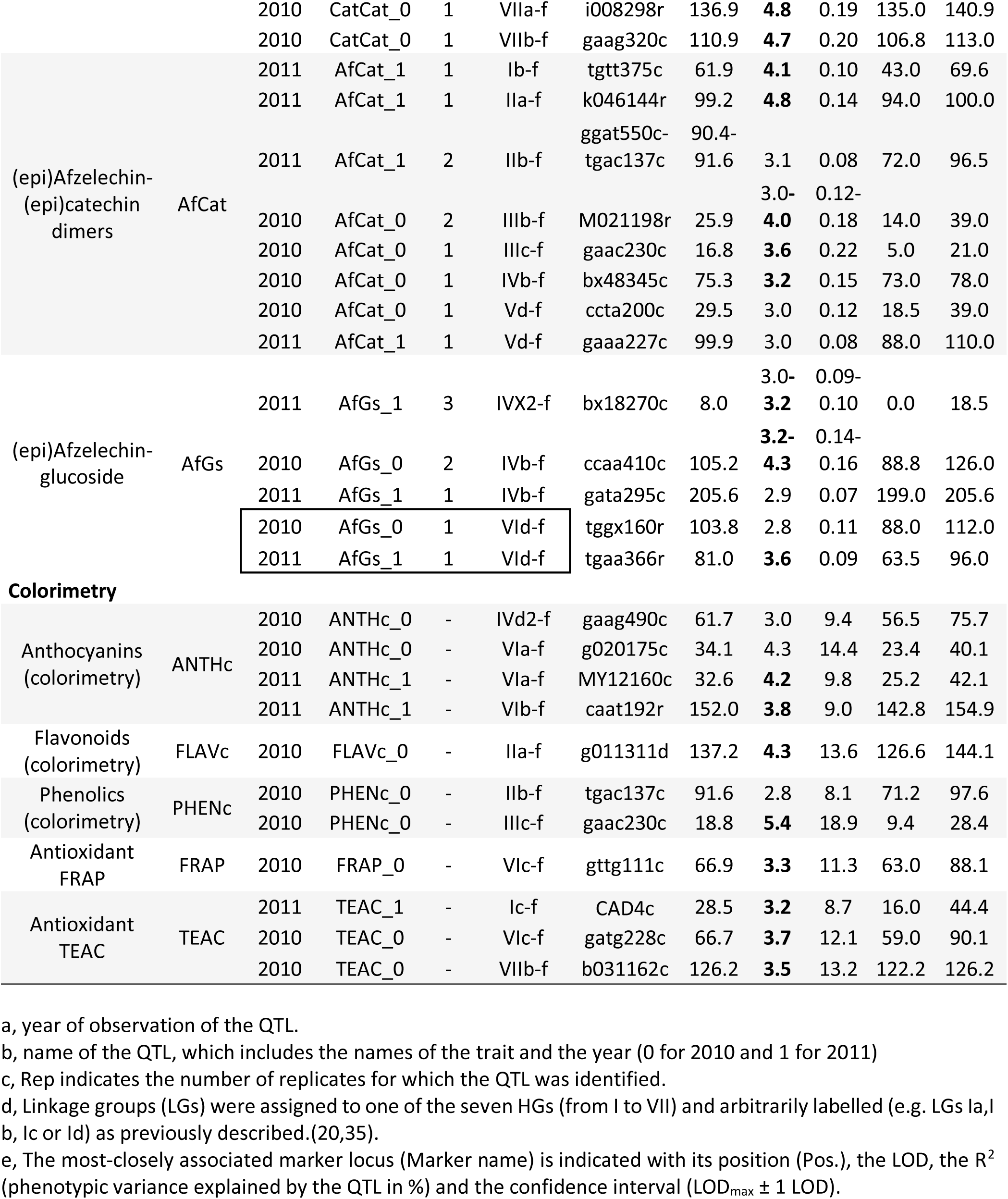
Significant QTLs detected for ‘Capitola’ (female). QTLs were identified for all the traits based on CIM analysis with LOD>LOD threshold (2.8 and 3.1 for respectively α = 0.10 or 0.05). QTLs identified in the two years of the study are in boxes. LOD in bold when significance level of the QTL is for α = 0.05.

**Supplemental Table 3.**
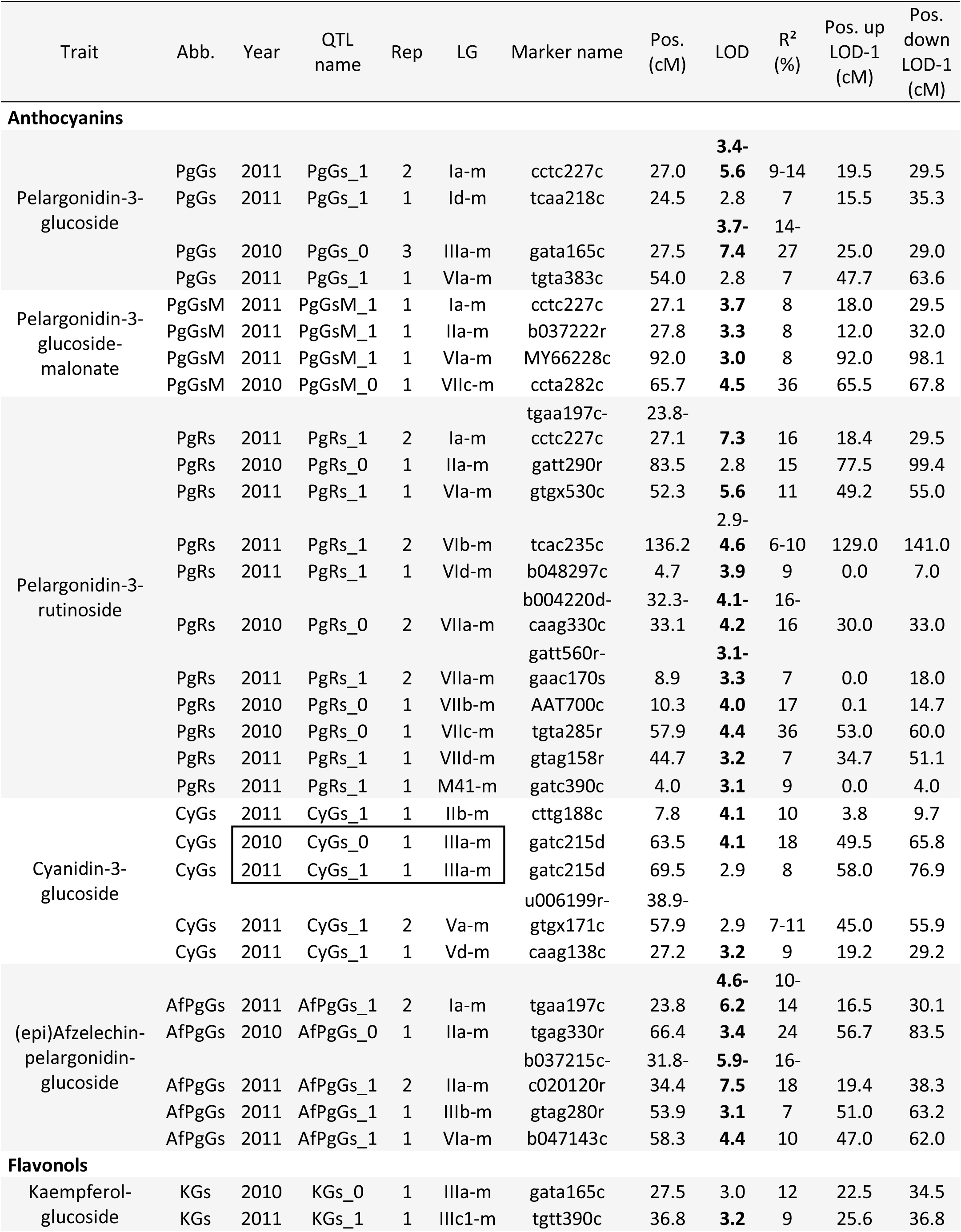
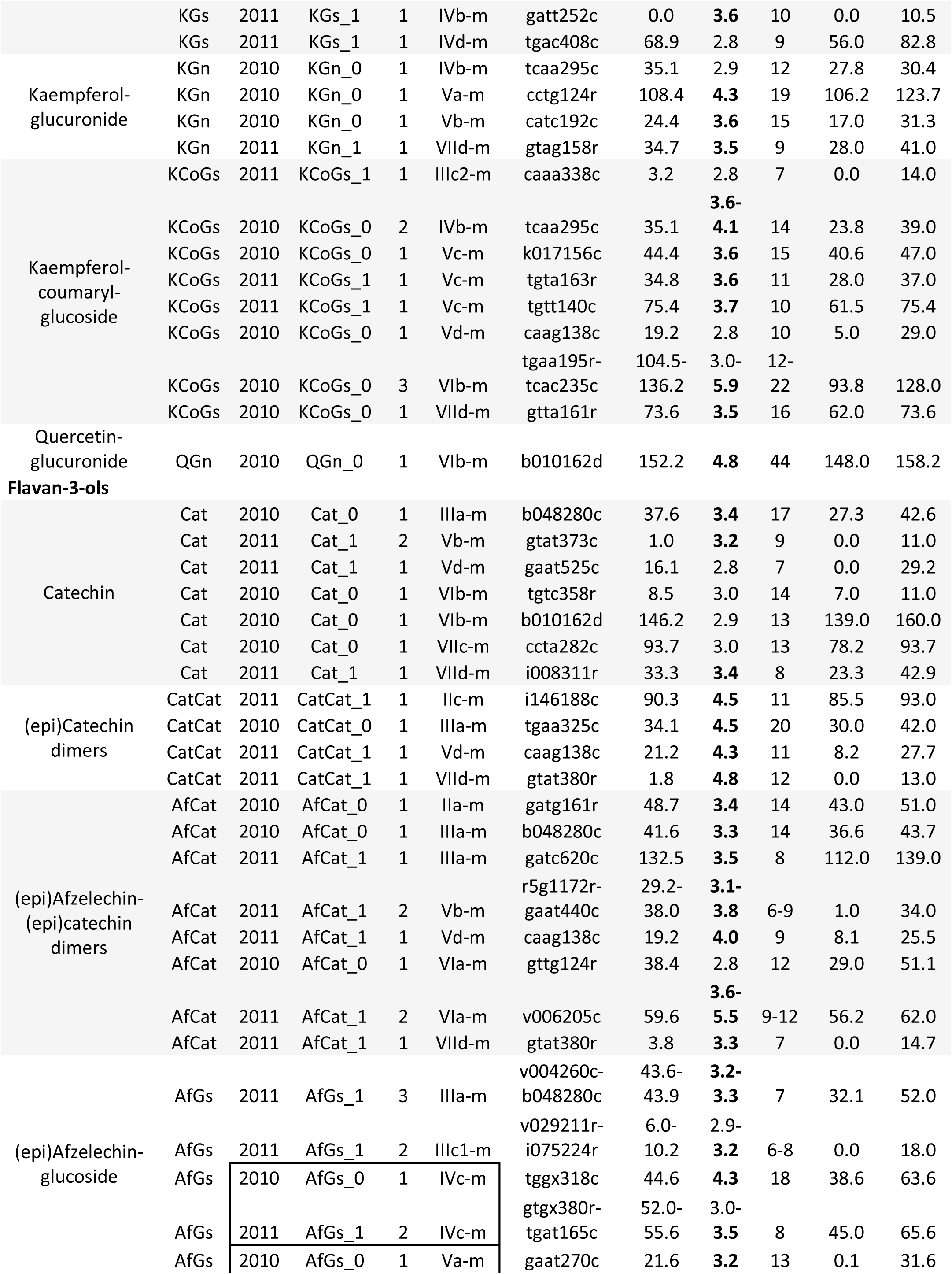
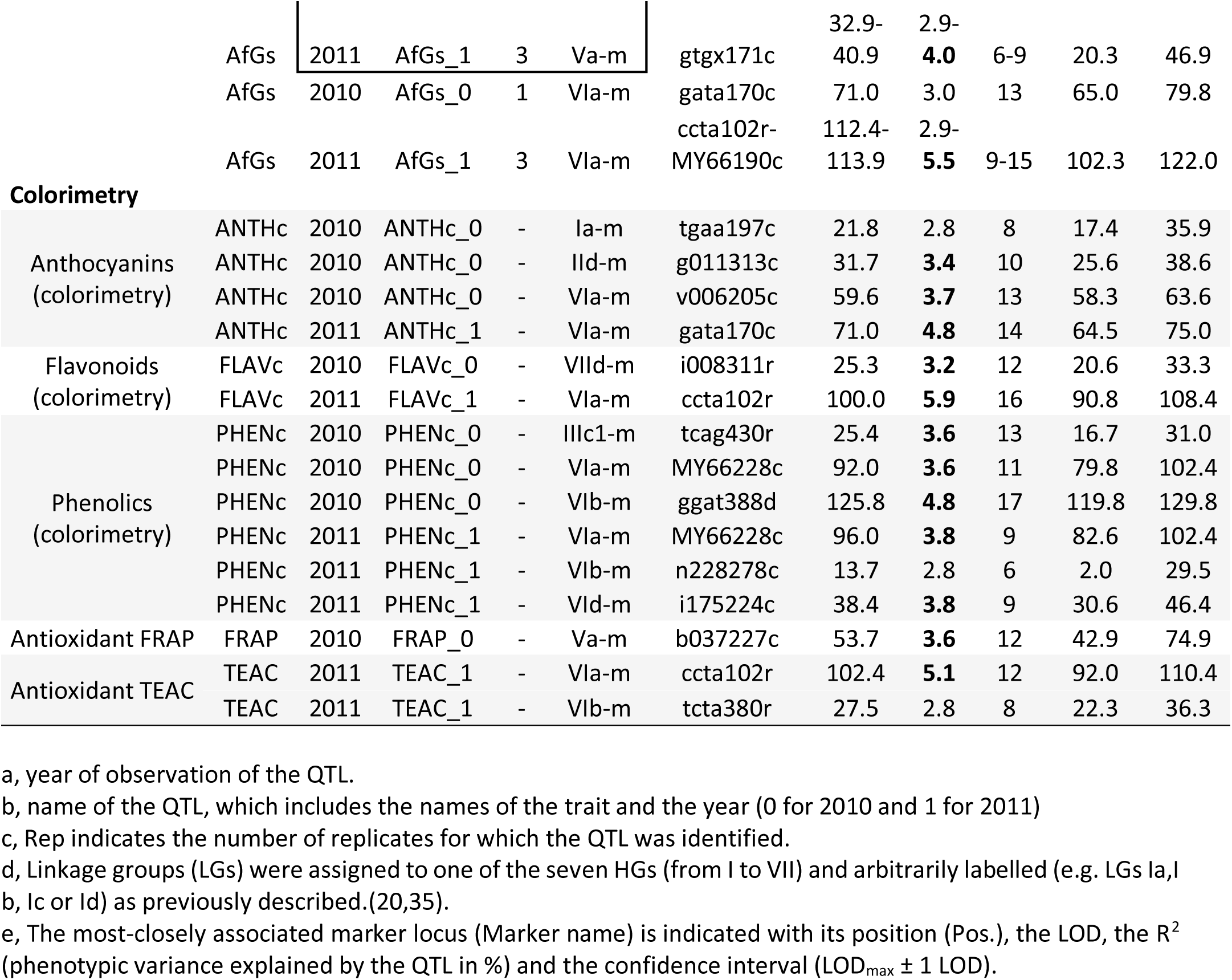
Significant QTLs detected for ‘CF1116’ (female). QTLs were identified for all the traits based on CIM analysis with LOD>LOD threshold (2.8 and 3.1 for respectively α = 0.10 or 0.05). QTLs identified in the two years of the study are in boxes LOD in bold when significance level of the QTL is for α = 0.05.

**Supplemental Table 4.**
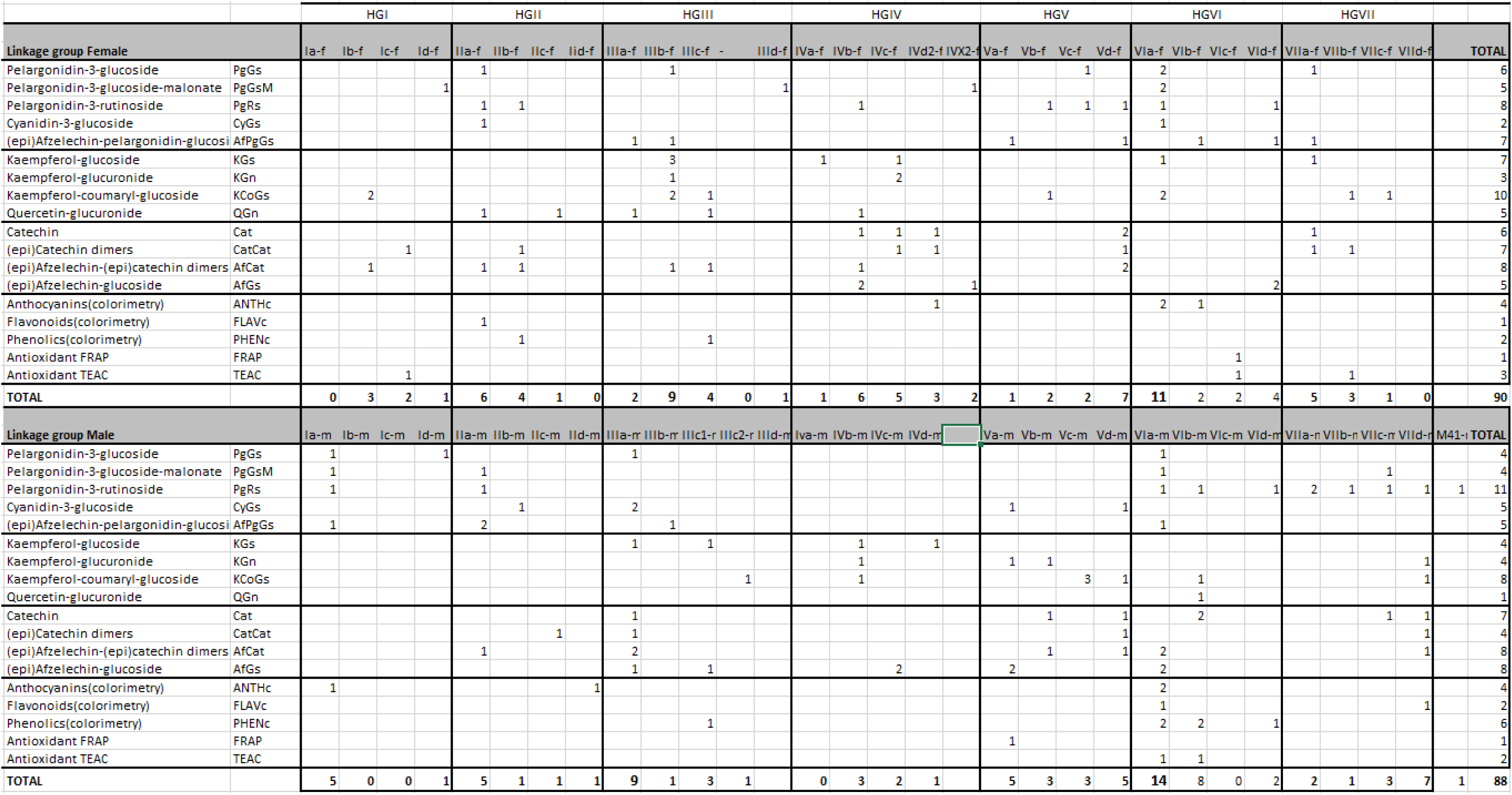
Distribution of the number of QTLs according to the homoeology groups and the linkage groups. Linkage groups (LGs) were assigned to one of the seven HGs and arbitrarily labelled (e.g. LG Ia, b, c or d) as previously described (Rousseau-Gueutin et al. 2008).

